# An improved habitat suitability index for the great crested newt

**DOI:** 10.1101/2022.12.15.520292

**Authors:** Emily Seccombe, Roberto Salguero-Gomez

## Abstract

Great crested newts (GCN) are a protected species whose conservation depends on the provision and protection of their breeding habitat. A habitat suitability index (HSI) developed in 2000 is extensively used in GCN conservation to assess breeding habitat quality. This two-decade old index is based on limited data and validation.

Here, we introduce a new HSI with an improved ability to reflect GCN presence/absence in UK ponds. This proposed HSI is easier, requires less data, and predicts GCN presence/absence better than the previous index.

To inform the new index, we used a dual approach to identify the relative importance of environmental criteria to predict GCN presence/absence. Firstly, we conducted a survey of 288 users of the GCN HSI to assess the perceived strengths and limitations of the existing index. Secondly, we analysed national datasets of GCN presence/absence and associated environmental data. Using these findings, we then tested various index modifications. The final modifications of the new HSI include: (i) using an arithmetic mean to combine variables, rather than a geometric mean, to reduce calculation errors and allow compensation between variables; (ii) excluding water quality and waterfowl impact as these lacked significant power to predict GCN presence/absence and were deemed inaccurate by HSI users; and (iii) changing the scoring relationship for pond area to better reflect current data and provide scores for ponds over 2000m^2^. Additionally, we present modified graphs to reduce errors in the calculation of pond density scores. We compared scores from the new and original HSIs using an independent dataset for validation, showing that the new HSI better reflects GCN present/absence (larger effect sizes and R-squared values) in comparison to the old HSI. Adopting this improved HSI will enable more effective conservation of the protected species via better-informed decision-making and monitoring.

## INTRODUCTION

Britain hosts populations of international importance for great crested newts (*Triturus cristatus*, Salamandridae – *GCN* hereafter, Dunford & Berry, 2013; Haysom et al., 2018). However, populations of this species are in serious decline (Beebee & Griffiths, 2000). The Joint Nature Conservation Committee’s GCN status report (2013-2018) concluded that there was an insufficient area and quality of occupied and unoccupied habitat for their long-term viability, and that habitat quality was decreasing (JNCC, 2019). Consequently, quantifying and identifying GCN habitat suitability is an important task for GCN conservation.

Due to their decline and international rarity, the GCN is protected under the Wildlife and Countryside Act (UK Government, 1981) and is a European Protected Species (The Council of the European Committees, 1992). Consequently, building developments that may harm GCN individuals or their habitat may require a licence, detailing mitigation requirements (English Nature, 2001). However, these mitigation measures are often suboptimally implemented and monitored (Edgar & Griffiths, 2004; Lewis et al., 2007). Due to the lack of conservation success and conflict with governmental housing development goals, Natural England launched a scheme in 2019 called District Level Licencing (Natural England, 2019). The scheme involves creating strategically-located replacement habitat to compensate for lost GCN habitat (Natural England, 2020). However, this approach relies on having sufficient understanding of the species’ habitat preferences to ensure the created habitat is suitable and to monitor the suitability of the habitat that is lost and created.

The GCN has narrower habitat requirements than other newt species in the UK (Wilkinson et al., 2011). Its aquatic habitat suitability depends on prey availability, presence of fish and waterfowl, number of nearby ponds, water quality, egg-laying material, pond desiccation frequency, and pond size (Beebee & Griffiths, 2000; Griffiths & Williams, 2000; Langton et al., 2000; Marklund et al., 2002; Skei et al., 2006; Denoël & Ficetola, 2008). Additionally, breeding sites need to be accessible from suitable terrestrial habitat that provides protection from predation, foraging opportunities, refuge from extreme weather, and hibernation opportunities (Gustafson et al., 2011).

Habitat suitability indices (*HSI*s hereafter) provide a process-based approach to model habitat suitability. For a HSI to be useful, it needs to accurately predict presence/absence of a given species based on limited data (Zajac et al., 2015). Importantly, while HSIs are widely used (e.g., Soniat et al., 2013, Bender et al., 1996), they are rarely based on peer-reviewed literature involving rigorous statistical analyses of environmental drivers of species presence/absence. Key concerns with HSIs include an overreliance on variable expert opinion (Johnson & Gillingham, 2004), as well as lack of uncertainty in estimates of presence/absence (Zajac et al. (2015), output validation (Brooks, 1997), and suitable frameworks for their objective evaluation (Roloff & Kernohan, 1999). Nonetheless, HSIs are often seen as a pragmatic solution for situations requiring management action (Brooks, 1997).

The current method for assessing a site’s suitability for GCNs is the HSI developed by Oldham et al. (2000). This index was created as “a simple model for use by the non-specialist, which provides conservationists with an informed view of the value of a site” (Oldham et al. 2000). In this HSI, scores for 10 criteria (Suitability Indices, *SI*s hereafter) are calculated and combined by taking their geometric mean, resulting in a single score ranging from zero (poor habitat) to one (suitable habitat) - see Table 1 and Oldham et al. (2000) and ARG UK (2010) for full details. The authors suggested that HSI scores would directly correlate with the carrying capacity of this species, though this link was never explicitly examined. Moreover, this HSI underwent limited validation - its parameterisation used only 72 ponds, predominantly located in only two of England’s 48 counties, but it is currently extrapolated across Britain, including as part of national reporting (Wilkinson & Arnell, 2013).

**Table 1:** Summary of suitability indices and their corresponding environmental criteria. This is a summary of the information provided in Oldham et al. (2001) and ARG UK (2010).

Oldham et al.’s (2000) HSI is used in a variety of key applications. These include site assessments for developments (Natural England, 2015) and assessment of mitigation success (Lewis et al., 2007, 2014). Importantly, the HSI is currently used in the UK’s statutory reporting on European Protected Species’ Favourable Condition Status (Reason, 2013). Furthermore, the HSI has been adapted for GCN monitoring schemes in continental Europe too (Jehle et al., 2011; Unglaub et al., 2015).

Despite the multiple applications of Oldham et al.’s (2000) HSI, there are significant concerns over its accuracy (Reason 2013; O’Brien et al. 2017; Buxton et al. 2021). Oldham et al. acknowledged the conjecture in the production of their HSI, noting that the index “can be upgraded easily as knowledge of crested newt habitat requirements improves”. Oldham’s HSI relies upon data from the 1991 National Amphibian Survey (Swan & Oldham, 1993; Oldham et al., 2000), now 30 years outdated. Despite advances in data availability since its creation, the HSI has remained largely unchanged. In 2007, the HSI was conservatively amended following a user-group workshop (ARG UK, 2010) to improve standardisation and usability. O’Brien et al. (2017) suggest a change to SI1 boundaries in Scotland.. Additionally, Buxton and Griffiths (2022) propose a change to how the HSI scores relate to categories. However, the index remains largely as created in 2000.

Only a few studies have assessed the HSI of the GCN. Reason (2013) found low repeatability of the index when calculating scores for 49 sites across England, concluding that the HSI is “clearly subjective”. Buxton et al. (2021) studied 146 ponds in north west England, finding significant proportions of ponds with GCN presence were categorised by the HSI as unsuitable habitat, and *vice versa*. Only waterfowl (SI6) and fish (SI7) indexes used in the HSI were significantly correlated with GCN presence/absence. Additionally, Lewis et al. (2007) tested the links between GCN count data and HSI scores, finding no significant relationship. O’Brien et al. (2017) found no significant relationship between GCN presence/absence and scores for pond area (SI2), waterfowl (SI6), fish (SI7) or pond density (SI8) at the northern edge of the species’ range, and proposed an improving zoning of the geographic location suitability index (SI1).

Motivated by the widespread application of Oldham’s HSI, here we develop an improved GCN HSI that increases its accuracy and simplicity in assessing the species’ habitat suitability. To do so, we surveyed HSI users and analysed national ecological datasets to identify the importance of different environmental factors in predicting GCN presence/absence across the UK. We then tested modifications and cross-validated the proposed HSI against the original index using an independent dataset of GCN presence/absence data. Our new HSI outperforms the original one as the goodness of fit is higher when using the new HSI compared to the original HSI, for both the original dataset and new independent validation dataset. The modified index uses the same underlying data as the original index, so where underlying data are available (e.g., pond area in m^2^ not just SI score), HSI scores can be recalculated with the new HSI, thus enabling comparison with existing data. We hope that the adoption of this HSI will allow more accurate estimation of habitat suitability for the GCN. The adoption of an improved HSI will enable more effective protection of this endangered species via better informed decision-making and monitoring.

## METHODS

We used a combination of methods to improve the accuracy and usability of the GCN habitat suitability index (HSI). We performed the statistical analyses with RStudio (v. 1.2.5001, R version 3.6.1, R Core Team, 2017) with the ‘tidyverse’ (Wickham et al., 2019) and ‘modeest’ (Poncet, 2019) packages for data preparation and description. The fully commented R script is available online (see Data Accessibility).

### User survey

We produced a survey targeted at users of Oldham’s HSI to assess perceptions of HSI recording accuracy, to elicit suggestions to improve the HSI, and to identify the extent of user support for the existing HSI. The target audience was users of the GCN HSI, aged over 18. Firstly, we conducted a pilot survey (Supp.Mat.1) to identify the required minimum sample size for the final survey, and to check the survey design viability. Following the pilot, the objectivity of the question wording was improved and the rating scales were modified to traditional Likert-like scales to reduce cognitive load (Chyung et al. 2017). The pilot revealed extensive answers to the optional qualitative questions, necessitating the use of textual analysis software, for which we chose NVivo (QSR International Pty Ltd., 2018). The final survey was published on SurveyMonkey and sent to 139 UK-based conservation organisations whose members are likely to use the HSI, such as ecology consultancies, biological record centres and conservation organisations. See Supp.Mat.2 for full survey text. The survey combined quantitative and qualitative questions. The quantitative questions asked for ratings of accuracy of records of the suitability indexes (SIs) and the HSI itself. 10 qualitative questions asked for information on limitations that reduced accuracy of SI records. Two further qualitative questions asked about additional factors for inclusion and other suggestions for improving the HSI.

The survey was completed by 288 respondents. The majority of respondents (86%) were professional ecologists, despite the variety of organisations contacted. As such, we could not test for an impact of respondent type on the answers. However, there was a wide range in survey participants’ years of experience, with a good level of response from experienced users (median of 14 years, ± 0.56 S.E.). Additionally, there was a reasonably high frequency of use of Oldham’s HSI amongst the respondents, with 50.4% using it 11 or more times per year. Neither years of experience (χ^2^_df=2_ = 1.386, *P* = 0.500) nor frequency of use of the index (χ^2^_df=4_ = 1.748, *P* = 0.782) significantly impacted scores of HSI accuracy.

We analysed the numeric questions through chi-square tests using the chisq.posthoc.test package (Ebbert, 2019). We applied content analysis in NVivo (QSR International Pty Ltd. 2018). to systematically examine the qualitative sections. We coded all answers, and subsequently joined codes together into themes based on frequency and similarity, following the practice set out in Bryman (2012). We used open coding rather than a pre-specified coding manual because of the exploratory nature of the survey (Bryman, 2012).

### Statistical analysis of ecological datasets

Next, we conducted a series of statistical analyses on ecological datasets of environmental correlates of GCN population presence/absence to better understand how environmental variables could be better integrated in an updated HSI. To this end, we sourced three datasets of sites with data on GCN presence/absence and HSI information. Cofnod supplied a dataset of 574 records from Wales collected between 2011 and 2019. The Amphibian and Reptile Conservation Trust supplied 852 records from the UK-wide National Amphibian and Reptile Recording Scheme (NARRS) project, 2013-2019. We also used two open-access datasets from Natural England, which include the eDNA research project with 5,866 records from 2017 to 2019 across England (Natural England Open Data, 2019), and the Evidence Enhancement Project with 3,137 records from six pilot areas across England, recorded in 2013 (Natural England Open Data, 2017). Additionally, an independent dataset from NatureSpace of 138 ponds for which GCN eDNA surveys were carried out in 2021 was used for verification of the new model. Prior to analysis, for each dataset, we performed data merging, homogenisation, and cleaning using the R packages ‘rnrfa’ (Vitolo et al., 2016), ‘EnvStats’ (Millard, 2019) and ‘lubridate’ (Grolemund & Wickham, 2011).

We performed a range of tests to identify statistically significant relationships between environmental variables and GCN presence/absence. We first fitted binomial regressions in a generalised linear model framework (R package ‘glmm’, Knudson, 2020) to test the relationship between the continuous variables (e.g., pond area, macrophyte cover) and GCN presence/absence (Skei et al., 2006). Next, we used Chi-square tests to test the relationship between categorical variables (e.g., waterfowl impact, fish presence) and GCN presence/absence. We used a Fisher’s test in place of Chi-square test for location due to imbalance data (most samples being from ‘Area A’). For those variables that were significantly correlated with GCN presence/absence, we then performed post-hoc tests of pairwise comparisons using the Bonferroni method (R package ‘chisq.posthoc.test’, Ebbert, 2019). We did not perform statistical analyses on GCN population abundance because the literature advises against using Oldham’s HSI in this context (Unglaub et al., 2015), and because the user survey findings showed that the HSI is mainly used to predict GCN presence/absence rather than abundance. Additional statistical details from the post-hoc tests are provided in Supp.Mat.3.

### Combining and testing modifications to the HSI

By combining the results of the survey and the statistical analyses regarding the environmental correlates of GCN presence/absence, we produced a list of candidate modifications to Oldham’s HSI. We tested some of these modifications where sufficient data were available to explore potential improvements in the HSI’s predictive ability. We then quantitatively assessed the potential improvement of the modifications on the HSI by adopting the approach implemented by Buxton et al. (2021). This involved comparing the distribution of GCN presence and absence amongst the five habitat suitability categories (Poor, Below Average, Average, Good, Excellent) using the new and modified HSIs. We then compared the resulting distributions of habitat suitability categories between Oldham’s HSI and our modified versions. We used Chi-square tests and the pertinent post-hoc tests of significant models on the frequency tables of HSI category and GCN presence/absence to identify if modified HSIs were more accurate in differentiating habitat quality than Oldham’s HSI. For these tests, we used the R packages ‘dplyr’ (Wickham et al., 2020), ‘EnvStats’ (Millard, 2019) and ‘forcats’ (Wickham, 2020) and ‘chisq.posthoc.test’ (Ebbert, 2019).

### Index cross-validation

To compare the original and new HSIs, we used the independent dataset (provided by NatureSpace, as noted above). The new HSI scores were calculated for each pond. We fitted binomial regressions in a generalised linear model framework (R package ‘glmm’, Knudson, 2020) to test the relationship between the HSI scores (new and original) and GCN presence/absence. Next, we used Chi-square tests and post-hoc tests to test the relationship between HSI category scores (Poor to Excellent) and GCN presence/absence, also using the Bonferroni method as above (R package ‘chisq.posthoc.test’, Ebbert, 2019).

## RESULTS

### Survey

The survey’s rating scale section (Supp.Mat.2) revealed much variation regarding the perceived accuracy of SIs (Figure 1). The chi-square tests showed that different SIs had significantly different perceived accuracy ratings (χ^2^_df=18_ = 77.454, P <0.001). Post-hoc tests showed that SI1 (geographic location) and SI9 (terrestrial habitat) had higher accuracy scores (P<0.001 and P<0.03 respectively), and that SI4 (water quality) had lower accuracy scores (P<0.001). The use of the HSI for GCN abundance estimations was rated significantly lower than for GCN presence/absence estimations (P<0.001; Figure S1). Overall, a quarter (26%) of respondents rated the HSI as ‘Very Inaccurate’ for estimating abundance.

**Figure 1:** The perceived accuracy of suitability index (SI) records for the great crested newt (GCN) by survey respondents varies across all 10 categories (Table 1). Bar charts show the frequency of our sampled 288 respondents selecting each accuracy rating option (from 1 to 7, where 1 is the lowest) in answer to the survey questions on the accuracy of records of SIs 1-10 (a-j).

Common themes arising from qualitative comments on the limitations of SI1-10 accuracy were identified. These themes included the subjectivity of SI measures (for SIs 2-10), the need to survey at certain times of year (particularly for SIs 2-7 and 10) and the difficulty interpreting SI graphs (particularly for SI2 (pond area) and SI8 (pond density). Another theme was criticism of surveyor skills, with key comments such as “Most GCN surveyors do not understand aquatic ecology”. Respondents also suggested a range of additional factors of potential importance for GCN habitat suitability that are not in the current HSI. The most frequent suggestions (with 17 comments each) were: known GCN populations in the area; chemical water tests; and disturbance/predation.

The respondents provided diverse suggestions for improvement of the HSI. These responses were grouped into themes which included: improving guidance to reduce subjectivity, making the HSI easier to use, using HSI results more appropriately and adapting the HSI to fit other circumstances (*e.g*., to apply to ditches). Many comments suggested misuse of the HSI, such as application outside of the recommended season and incorrect geometric mean calculation. Hierarchy plots of the responses to each qualitative question, with further detail and quantification, are given in Supp.Mat.4.

### Ecological determinants of GCN presence

Scores of continuous SIs were significantly higher in sites with GCNs present than sites with GCNs absent for SI5 (shade), SI8 (pond density), and SI10 (macrophytes) (all P<0.001; Figure 2). Binomial regressions found no significant difference in SI2 (pond area) scores for ponds with GCNs present or absent.

**Figure 2:** Minimal differences in scores of continuous suitability indexes (SIs) are apparent between sites with great crested newt (GCN) absent (salmon) or present (green). Box and whisker plots show (a) SI2: pond area, (b) SI5: shade, (c) SI8: pond count, and (d) SI10: macrophyte scores for sites with GCNs present or absent. “n.s.” indicates statistically non-significant differences between SI scores in ponds with GCN absent or present; “**” indicates statistically significant differences at P<0.001 between SI scores in ponds with GCN absent or present.

Of the categorical SIs, significant differences in scores at ponds with GCNs present or absent were apparent for SI3 (pond permanence), SI4 (water quality), SI6 (waterfowl), SI7 (fish), and SI9 (terrestrial habitat) (P<0.001, for full residuals, please see Supp.Mat.3a) - see Figure 3. The post-hoc tests showed that these differences were in the directions expected (i.e., higher SI values correlated with presence of the species) for most of these SIs, except for SI6 and SI7. Surprisingly, the post-hoc test for SI6 showed that ponds with GCNs absent had significantly more scores of 1 (indicating waterfowl absence) and fewer 0.67 scores (indicating minor waterfowl impact) and vice versa for ponds with GCNs present (P<0.001). The post-hoc test for SI7 also showed an unexpected pattern: ponds with GCNs absent had more 0.01 scores (indicating major fish impact), more 1 scores (indicating fish absence) and fewer 0.67 scores (indicating possible fish impact) and *vice versa* (P<0.001). No significant difference was found for SI1 scores (geographic location) at ponds with GCNs present than ponds with the species recorded as absent.

**Figure 3:** Most categorical suitability indexes (SIs; Table 1) for the great crested newt (GCN) show marginally higher frequency of ponds with GCN present (green) for higher score categories, and lower frequency of ponds with GCN absent (salmon) for lower score categories. This pattern is not apparent for waterfowl (SI6), and is hard to assess for pond location (SI1) due to the predominance of a score of 1, meaning few samples were from the edges of GCN distribution. Bar charts show the frequency of score levels for sites with GCNs present or absent for (a) SI1: geographic location, (b) SI3: pond permanence, (c) SI4: water quality, (d) SI6: waterfowl, (e) SI7: fish, and (f) SI9: terrestrial habitat. “n.s.” indicates statistically non-significant differences between SI scores in ponds with GCN absent or present, “**” indicates the statistically significant differences at P<0.001 between SI scores in ponds with GCN absent or present.

Binomial regression found a significant relationship (slope=0.703, d.f.=9,046, P<0.001) between HSI scores and presence/absence of GCNs (Figure 4). We found significant differences in the HSI category frequencies for ponds with GCNs present or absent (χ^2^_df=4_=245.04, P<0.001). As expected, post-hoc tests showed that ponds without GCNs had more Below Average and Poor ponds and fewer Excellent and Good ponds than ponds with GCNs (P<0.001 in all cases). This reflects the patterns shown in Figure 4a.

**Figure 4:** Habitat suitability index (HSI) scores for the great crested newt (GCN) are higher in ponds where the species is present. HSI (a) categories and (b) scores are shown for sites with (salmon) or without (green) GCNs. (a) The bar chart shows higher frequencies of ponds with GCN present and lower frequencies of ponds with GCN present for the higher HSI categories (Average, Good, and Excellent). The panel also shows lower frequencies of ponds with GCN present and higher frequencies of ponds with GCN present for the lower HSI categories (Poor and Below average). (b) Box and whisker plot of scores showing higher HSI scores for sites where GCNs are present.

### HSI improvements

Informed by the previous results, a list of potential modifications is presented in Tables 2 and 3. Some of these modifications require additional data or involve changes to the written guidance. The modifications that could be tested with the data available for this study were explored and results present below. Additionally, new graphs were produced to aid reading of continuous SIs (see Figures S4 - S6), and formulae provided in Supp.Mat.5.

**Table 2:** Potential modifications to improve the great crested newt (GCN) suitability indices (SIs) within the habitat suitability index (HSI), inferred from peer-review literature, user survey results, and ecological data analysis performed in this research.

**Table 3:** Potential modifications to improve the great crested newt (GCN) the habitat suitability index (HSI), inferred from peer-review literature, user survey results, and ecological data analysis performed in this research.

Two of the datasets were found to have incorrect HSI calculations due to errors in geometric mean calculation. To overcome this issue, new HSI scores were calculated using the arithmetic mean. The distribution of HSI score categories remained significantly different between sites with GCN present or absent (χ^2^_df=4_=261.85, P<0.001), see Supp.Mat.3b for residuals from the post-hoc test. Therefore, the arithmetic mean-based HSI is comparable in terms of reflecting GCN presence/absence, but avoids the likelihood of calculation mistakes of the geometric mean.

SI8 (pond density) scores were not calculated correctly in all the datasets (in the NARRS dataset, the SI8 scores were not divided by π, resulting in no sites scoring between 0.1 and 0.7), a problem also noted by survey respondents. To prevent these errors, a modified graph was produced (Figure S5) that avoids the need for division by *π*.

New HSI scores were created by excluding the SIs that received low accuracy ratings in the survey. Independent exclusion of SI4 (water quality) and SI6 (waterfowl) were found to be beneficial (see Supp.Mat.3c and 3d)., so these SIs were then excluded simultaneously. The distribution of sites between HSI categories remained significantly different for ponds with GCN present or absent (χ^2^_df=4_=369.83, P<0.001). The post-hoc test showed the modified HSI was better than the original (larger residuals) for distinguishing Excellent and Poor ponds, and comparable for other categories (Supp.Mat.3e).

A new scoring method for SI2 (pond area) was created by plotting new data to create a new scoring graph (see Figure S2 and S3). HSI scores were recalculated with the new SI2 scores. The distribution of sites between HSI categories remained significantly different for ponds with GCN present or absent with the new SI2 scores (χ^2^_df=4_=297.51, P<0.001). The post-hoc test showed the modified HSI was better at distinguishing between ponds in the Below Average, Excellent, and Good categories, although very slightly worse for the Poor category (see residuals in Supp.Mat.3f) in comparison to the original HSI.

The modifications detailed above were then combined and new HSI scores were calculated using these modifications. The scores were grouped in categories with new boundaries to create more equal splits to facilitate interpretation (Table 4). The new scores and category frequencies are shown in Figure 5, which can be compared to the original HSI scores and category frequencies in Figure 4.

**Figure 5:** Distribution of (a) HSI categories and (b) HSI scores for sites with great crested newts (GCNs) present or absent under the newly proposed habitat suitability index (HSI). HSI scores are statistically higher in ponds where GCNs are present (P<0.001) - see Supp.Mat.3g. (a) Bar chart showing higher frequencies of ponds with GCN present and lower frequencies of ponds with GCN present for the higher HSI categories (Average, Good and Excellent). This panel shows lower frequencies of ponds with GCN present and higher frequencies of ponds with GCN absent for the lower HSI categories (Poor, Below average). (b) Box and whisker plot of the newly proposed HSI scores showing higher HSI scores for sites with where GCNs are present (P<0.001) - see Supp.Mat.3h. The interquartile range of modified HSI scores in Figure 5.B is higher and more compact than that of Oldham’s HSI shown in Figure 4.B, necessitating the new category divisions proposed in Table 4.

**Table 4:** Habitat suitability index (HSI) categories (left column) and corresponding values (centre column) under the original habitat suitability index (ARG UK, 2010) and with the new modified habitat suitability index (right column).

The chi-square test of the new HSI categories and GCN presence/absence showed strong significance (χ^2^_df=4_= 533.11, P<0.001). The residuals for all categories are larger than for the original HSI (see Supp.Mat.3g for residuals).

#### Cross-validation

Both the original and new HSIs lacked predictive power for GCN presence/absence at high significance values with the independent dataset. However, the new HSI had higher effect sizes and lower residuals from the binomial regression than the original HSI scores (see Figure 6, and Supp.Mat.3h for full residuals). Following chi-square tests to look at HSI category scores, post-hoc tests were carried out, but no significant differences in GCN presence/absence between HSI categories were found (see Supp.Mat.3i).

**Figure 6.** The newly proposed HSI outperforms the old HSI both in terms of effect sizes and goodness of fit when predicting presence/absence of the great created newt (GCN). Statistical results of the binomial regression testing the relationship between HSI scores and GCN presence/absence. The length of the arrows is proportional to the effect size for the output of the binomial regression, under the two HSIs, first tests on the original dataset of the paper, and secondly on the independent dataset used for verification. The circles show the R2 values - with the new HSI, these values are larger (more so for the original dataset) suggesting a better goodness-of-fit between the new HSI and the GCN presence/absence data. The effect sizes are only statistically significant when using the larger, original dataset.

## DISCUSSION

We examined potential improvements to existing habitat suitability index (HSI) for the great crested newt (GCN) proposed by Oldham et al. (2000), as adapted by ARG UK (2010). Through a combined approach using a HSI-user survey and robust statistical analyses of GCN and environmental data across the UK, we found that the HSI could be improved by removing the indices for SI4 (water quality) and SI6 (waterfowl), combining the index values with an arithmetic rather than geometric mean, and creating new scoring scheme for SI2 (pond area). Thus, we introduce an improved HSI for GCN in the UK.

This study identified a range of limitations of the original HSI, including limitations to SI recording accuracy identified from the user survey, with common themes including subjectivity, surveyor skill, and temporal variability. The findings echo wider literature about HSIs, such as concerns about a lack of objectivity (Brooks, 1997). However, HSIs are intended as applicable management tools and thus their accuracy will pragmatically always be limited (Johnson & Gillingham, 2004). Nonetheless, the limitations to the HSI given in the survey and the criticism in the existing literature justify further work to improve the HSI, particularly given its conservation importance for the species.

Concern over application of the HSI to reflect the GCN abundance was a key theme arising from the survey. These findings corroborate concerns raised by Lewis et al. (2007) and Buxton et al. (2021). The HSI guidance (NARRS, n.d., and ARG UK, 2010) and Natural England’s GCN webpage (Natural England, 2015b) specify that the HSI should not be used as replacement for population surveys or to predict abundance, yet this practice continues.

The ecological data analysis suggested that the original HSI has limited ability to differentiate GCN suitability as inferred from presence/absence. Our results showed that many ponds in low HSI categories (‘Poor’ or ‘Below Average’) have GCNs present, and many ponds in high HSI categories (‘Good or ‘Excellent’) have GCNs recorded as absent. This finding echoes the results of Buxton et al. (2021) at a national stage. This demonstrates the need to continue investigating HSIs after their creation - such research is lacking for other species’ HSIs too, as highlighted in Brooks (1997).

A further limitation of the HSI identified by the ecological analysis is that only some SIs were significantly correlated with GCN presence/absence. SI1 (location) and SI2 (pond size) were not significantly correlated with GCN presence/absence. The latter finding corroborates the findings of Denoël and Ficetola (2008). SI5, SI8, SI9, and SI10 (shade, pond density, terrestrial habitat quality, and macrophytes) scores were positively correlated with GCN presence. The significant relationship for SI8 contradicts the lack of effect of pond density in Denoël and Ficetola’s study (2008). SI3 (pond permanence), SI4 (water quality), SI6 (waterfowl) and SI7 (fish) had mixed results. The lack of clear correlations for SI6 and SI7 reflect the conflicting literature (Marklund et al., 2002, Skei et al., 2011, and Denoël and Ficetola, 2008). The findings of SI significance outlined above contradict regional studies of the HSI, suggesting that regional variation is important and supports the need for national studies into the HSI.

This study also identified errors in HSI calculation. These errors include incorrectly calculating pond density scores (as in the NARRS dataset) or not correctly adjusting the geometric mean calculation for the number of SIs recorded (as in the NARRS dataset and the Natural England eDNA dataset). This finding adds to the need for increased validation of HSIs (Brooks, 1997), to ensure that not only does the HSI provide an accurate reflection of the species’ habitat requirements, but that the HSI is easy to calculate correctly.

This study produced a range of potential modifications to improve the HSI (summarised in Tables 2 and 3). Those modifications which could be tested with the available data led to a new proposed HSI with the following modifications: an arithmetic mean is used to combine SIs instead of a geometric mean; a new SI2 (pond size) scoring relationship is used; SI4 (water quality) and SI6 (waterfowl) are removed; and SI8 (pond density) scores are correctly calculated, aided by a modified graph. Following the approach of Buxton et al. (2021), the distribution of ponds with GCNs present or absent between different HSI categories was used to assess HSI accuracy. According to this method, the new HSI is better than the original at distinguishing GCN habitat suitability, when assuming that suitability can be inferred from presence/absence (Lewis et al., 2007). The new HSI also has the advantage of having fewer ponds in the Average category which was difficult to interpret. It should be easier to conduct due to simpler calculation and having fewer SIs. The exclusion of SI4 (water quality) and SI6 (waterfowl) avoids the problems in assessing water quality and waterfowl impact noted by survey respondents. The new graphs (Figures S4-S6) and formulae (Supp.Mat.5) also support ease-of-use of the new HSI. The proposed HSI can be applied to existing data, as long as there is information on the underlying environmental variables (e.g., pond size, not just SI2 score) so that new HSI scores can be calculated for prior records, allowing comparison of HSI scores over time.

The results of the cross-validation binomial regression using generalised linear models suggests that the new HSI scores better reflect GCN presence/absence. Both indexes are limited in predictive power, although the new index has higher effect sizes and R^2^ values, whilst importantly using fewer variables. The lack of significance when using either HSI on the verification dataset may be due to the small sample size, or irregularities of the data, perhaps due to all the data being from a single year.

The complex nature of the ecological data used introduces limitations, such as non-random site selection. Another limitation comes from the low detection probabilities for GCNs (Sewell et al., 2010). Spatial autocorrelation is a further risk in this data (Crase et al., 2012). One particular difficulty comes from the inevitably false assumption that populations are in equilibrium with the environment, thereby creating an issue with using presence/absence as a proxy for habitat suitability (Latimer et al., 2006). This assumption is particularly problematic for species with metapopulation dynamics (Kupfer & Kneitz, 2000).

This research provides a case study for updating HSIs. As highlighted by Brooks (1997), there is a lack of research into HSIs and they are rarely modified or improved over time, despite high management importance and increased data availability.

Potential additional modifications could be further investigated based on this study. Additional research could inform weightings for the SIs (following Ray & Burgman, 2006) or could provide uncertainty estimates of HSI output (following the work of Bender et al., 1996; Burgman et al., 2001; Zajac et al., 2012). The research presented here provides an evidence-based starting point for discussions on how to best adapt the HSI and the accompanying guidance for users.

## CONCLUSION

Our multidisciplinary approach, which combines expert GCN elicitation together with robust environmental statistical analyses, shows tangible ways to improve the existing GCN HSI. We have presented greater insight into the current HSI’s limitations and have proposed a modified HSI which better reflects likely GCN presence/absence. We also presented suggestions of further potential modifications warranting additional research. Our proposed new HSI has three key modifications: (i) using an arithmetic (rather than geometric) mean to combine variables, to reduce calculation errors and allow compensation between variables; (ii) excluding water quality and waterfowl impact as these lacked significant power to predict GCN presence/absence and were deemed inaccurate by HSI users; and (iii) changing the scoring relationship for pond area to better reflect current data and provide scores for ponds over 2000m^2^. We argue that the improved HSI developed in this study is better placed to inform more accurate assessment of the habitat suitability of ponds for GCNs, and thereby provide more accurate monitoring of habitat trends and guide GCN conservation and development-mitigation. This research is important due to the extensive use of the HSI in GCN conservation. and we hope that a new HSI will ultimately be implemented as a result of this research.

## Supporting information

Cover Page

## ACKNOWLEDGEMENTS

We thank Dr P. Berry (University of Oxford) and Dr J. Wilkinson (Amphibian and Reptile Conservation Trust) for their advice. We thank the Amphibian and Reptile Conservation Trust, Cofnod, and NatureSpace for permitting access to their data, and the survey participants. Author 2 was supported by a NERC Independent Research Fellowship (NE/M018458/1).

## DATA ACCESSIBILITY

The data, code, and supplementary materials can be found here: https://osf.io/2yt6r/

The Natural England data can be found here: https://naturalengland-defra.opendata.arcgis.com/ - see references for Natural England Open Data 2017 and 2019. Data uploaded 22.03.2017.

