## Supplementary Materials for "An improved habitat suitability index for the great crested newt"

**SUPPLEMENTARY MATERIALS AND FIGURES:**

**Contents**

- **Supplementary Materials 1: Pilot survey**
- **Supplementary Materials 2: Final survey text**
- **Supplementary Materials 3: Additional statistical information from post-hoc tests**
- **Supplementary Materials 4: Qualitative survey results**
- **Supplementary Materials 5: Line formulae for new graphs**

**Supp. Mat. 1 Pilot survey text**

Great Crested Newt Habitat Suitability Index Users Survey PILOT

#### General Information

**We appreciate your interest in participating in this online survey of users of the Great Crested Newt (GCN) Habitat Suitability Index (HSI).**

**Please read through this information before agreeing to participate.**

**You may ask any questions before deciding to take part by contacting the researcher (details below).**

**This survey aims to collect data on the perceived accuracy of the HSI as a tool for assessing the habitat suitability of ponds for GCN. This information will be used alongside ecological data to build a new HSI that is easy-to-use and accurate.**

**You will be given 27 questions to answer. All questions are optional. This should take about 15 minutes (PILOT ONLY: Please record the time taken to complete the survey to answer the final question). Whilst you must have used the GCN HSI (either as a volunteer or professional), no further knowledge or experience is required. The HSI referred to throughout is that made by Oldham et al. (2000) and can be found here: https://www.arguk.org/get-involved/projects-surveys/great-crested-newt-habitat-suitability-index**

**Please note that your participation is voluntary. If you do decide to take part, you may withdraw at any point for any reason before submitting your answers by closing the browser.**

**How will my data be used?**

**Your answers will be completely anonymous, and we will take all reasonable measures to ensure that they remain confidential. Your data will be securely stored in a password-protected file until it has been checked for personal data. The results may be used in academic publications. Your IP address will not be stored. No questions ask for personal or sensitive information. Please do not enter personal information as answers to the questions. Research data will be stored for the duration of the study period (until 1st September 2020).**

**Who will have access to my data?**

**SurveyMonkey is the data controller with respect to your personal data and, as such, will determine how your personal data is used. Please see their privacy notice here: https://www.surveymonkey.com/mp/legal/privacy-policy/. SurveyMonkey will share only fully anonymised data with the University of Oxford, for the purposes of research. Results will be shared with the Amphibian and Reptile Conservation Trust.**

**We would also like your permission to use your anonymised data in future studies, and to share data with other researchers (e.g. in online databases). Any personal information that could identify you will be removed or changed before files are shared with other researchers or results are made**

**public.**

**Responsible members of the University of Oxford and funders may be given access to data for monitoring and/or audit of the study to ensure we are complying with guidelines, or as otherwise required by law.**

**This survey is for an MSc project. The Principal Researcher is [redacted], who is attached to the School of Geography and the Environment at the University of Oxford. This project is being completed under the supervision of [redacted].**

**This project has been reviewed by, and received ethics clearance through the University of Oxford Central University Research Ethics Committee (reference: SOGE 1A2020-66).**

Who do I contact if I have a concern about the study or I wish to complain?
If you have a concern about any aspect of this study, please speak to [redacted] or their supervisor [redacted] and we will do our best to answer your query. We will acknowledge your concern within 10 working days and give you an indication of how it will be dealt with. If you remain unhappy or wish to make a formal complaint, please contact the Chair of the Research Ethics Committee who will seek to resolve the matter as soon as possible:
School of Geography and the Environment Departmental Research Ethics Committee Chair: Prof Jim Hall, Contactable via Gillian Willis, School of Geography and the Engironment, University of Oxford, South Parks Road, Oxford, OX1 3QY, UK.

* 1. Please note that you may only participate in this survey if you are 18 years of age or over. If you are not, please exit the survey.

- ◯ Yes - I certify that I am 18 years of age or over

* 2. If you have read the information above and agree to participate with the understanding that the data (including any personal data) you submit will be processed accordingly, please check the relevant box below to get started. If not, please exit the survey.

- ◯ Yes - I agree to take part

UNIVERSITY OF OXFORD

#### Great Crested Newt Habitat Suitability Index Users Survey PILOT

##### Part 1 of 3: Your use of the Habitat Suitability Index

1. In what capacity do you use the HSI for GCNs? (If more than one category applies, please select the capacity under which you usually use the index)

- ◯ Volunteer – independent
- ◯ Volunteer - ecological consultancy or similar
- ◯ Volunteer - conservation organisation
- ◯ Other (please specify)
- ◯ Professional - conservation organisation
- ◯ Professional - ecological consultancy or similar

2. For approximately how many years have you used the HSI for GCNs?

0 21

3. Approximately how many times per year do you use the HSI for GCNs (excluding years when not doing any)?

- ◯ 1-10
- ◯ 11-50
- ◯ >50

4. How accurate do you think the HSI is for predicting GCN presence/absence?
[10 is high and 0 is low]

0 10

5. How accurate do you think the HSI is for predicting GCN abundance?
[10 is high and 0 is low]

0 10

UNIVERSITY OF OXFORD

### Great Crested Newt Habitat Suitability Index Users Survey PILOT

#### Part 2 of 3: Factors in the HSI

You may find it helpful to refer to an HSI guidance document when answering the following questions, such as the document available here: http://www.narrs.org.uk/documents/HSI%20guidance.pdf

The first factor in the HSI is Geographic Location (SI1). Sites are scored according to the zone in which they occur, using a figure of the UK.

1. How accurate do you estimate your own records of SI1 to be?
[10 is high for complete accuracy, and 0 is low for being no better than random guessing]

0 10

2. What limits the accuracy of your records of SI1? (Optional)

The second factor is pond area (SI2). The index value is read off a graph.

3. How accurate do you estimate your own records of SI2 to be?
[10 is high for complete accuracy, and 0 is low for being no better than random guessing]

0 10

4. What limits the accuracy of your records of SI2? (Optional)

The third factor is pond permanence (SI3). The index value is given by one of four categories (never, rarely, sometimes or annually dries out)

5. How accurate do you estimate your own records of SI3 to be?
[10 is high for complete accuracy, and 0 is low for being no better than random guessing]

0 10

6. What limits the accuracy of your records of SI3? (Optional)

The fourth factor is water quality (SI4). The index value is given by one of four categories (good, moderate, poor or bad).

7. How accurate do you estimate your own records of SI4 to be?
[10 is high for complete accuracy, and 0 is low for being no better than random guessing]

0 10

8. What limits the accuracy of your records of SI4? (Optional)

The fifth factor is shade (SI5). The index value is read off a graph, based on the percentage of the pond perimeter which is shaded.

9. How accurate do you estimate your own records of SI5 to be?
[10 is high for complete accuracy, and 0 is low for being no better than random guessing]

0 10

10. What limits the accuracy of your records of SI5? (Optional)

The sixth factor is the impact of waterfowl (SI6). The index value is given by one of three categories (absent, minor, major).

11. How accurate do you estimate your own records of SI6 to be?
[10 is high for complete accuracy, and 0 is low for being no better than random guessing]

0 10

12. What limits the accuracy of your records of SI6? (Optional)

The seventh factor is the impact of fish (SI7). The index value is given by one of four categories (absent, possible, minor, major).

13. How accurate do you estimate your own records of SI7 to be?
[10 is high for complete accuracy, and 0 is low for being no better than random guessing]

0 10

14. What limits the accuracy of your records of SI7? (Optional)

The eighth factor is pond count (SI8). The index value is read off a graph, based on the number of ponds within 1km.

15. How accurate do you estimate your own records of SI8 to be?
[10 is high for complete accuracy, and 0 is low for being no better than random guessing]

0 10

16. What limits the accuracy of your records of SI8? (Optional)

The ninth factor is terrestrial habitat (SI9). The index value is given by one of four categories (good, moderate, poor, none).

17. How accurate do you estimate your own records of SI9 to be?
[10 is high for complete accuracy, and 0 is low for being no better than random guessing]

0 10

18. What limits the accuracy of your records of SI9? (Optional)

The tenth factor is macrophyte cover (SI10). The index value is read off a graph

19. How accurate do you estimate your own records of SI10 to be?
[10 is high for complete accuracy, and 0 is low for being no better than random guessing]

0 10

20. What limits the accuracy of your records of SI10? (Optional)

#### Great Crested Newt Habitat Suitability Index Users Survey PILOT

##### Part 3 of 3: Other factors

1. Do you think there are other environmental factors that may impact the habitat suitability of ponds for Great Crested Newts that are not currently included in the HSI? If so, please list below.

2. Do you have other suggestions for the improvement of the HSI for GCN? If so, please list below.

3. PILOT ONLY: How long did you spend completing the survey?

4. PILOT ONLY: Do you have any comments as pilot participants relating to the survey design?

**Supp. Mat. 2 Final survey text**

Great Crested Newt Habitat Suitability Index Users Survey

#### General Information

**We appreciate your interest in participating in this online survey of users of the Great Crested Newt (GCN) Habitat Suitability Index (HSI).**

**Please read through this information before agreeing to participate.**

**You may ask any questions before deciding to take part by contacting the researcher (details below).**

**This survey aims to collect data on the perceived accuracy of the HSI as a tool for assessing the habitat suitability of ponds for GCNs. This information will be used alongside ecological data to investigate potential improvements to the HSI.**

**You will be given 27 questions to answer. All questions are optional. This should take about 10 minutes. Whilst you must have used the GCN HSI (either as a volunteer or professional), no further knowledge or experience is required. The HSI referred to throughout is that made by Oldham et al. (2000) and updated in the ARG UK Advice Note 5 guidance which can be found here: https://www.arguk.org/get-involved/projects-surveys/great-crested-newt-habitat-suitability-index**

**Please note that your participation is voluntary. If you do decide to take part, you may withdraw at any point for any reason before submitting your answers by closing the browser window.**

**How will my data be used?**

**Your answers will be completely anonymous, and we will take all reasonable measures to ensure that they remain confidential. Your data will be securely stored in a password-protected file until it has been checked for personal data. The results may be used in academic publications. Your IP address will not be stored. No questions ask for personal or sensitive information. Please do not enter personal information as answers to the questions. Research data will be stored for the duration of the study period (until 1st September 2020).**

**Who will have access to my data?**

**SurveyMonkey is the data controller with respect to your personal data and, as such, will determine how your personal data is used. Please see their privacy notice here: https://www.surveymonkey.com/mp/legal/privacy-policy/. SurveyMonkey will share only fully anonymised data with the University of Oxford, for the purposes of research. Results will be shared with the Amphibian and Reptile Conservation Trust.**

**We would also like your permission to use your anonymised data in future studies, and to share data with other researchers (e.g. in online databases). Any personal information that could identify you will**

**be removed or changed before files are shared with other researchers or results are made public.**

**Responsible members of the University of Oxford and funders may be given access to data for monitoring and/or audit of the study to ensure we are complying with guidelines, or as otherwise required by law.**

**This survey is for an MSc project. The Principal Researcher is [redacted], who is attached to the School of Geography and the Environment at the University of Oxford. This project is being completed under the supervision of [redacted]**

**This project has been reviewed by, and received ethics clearance through the University of Oxford Central University Research Ethics Committee (reference: SOGE 1A2020-66).**

<u>Who do I contact if I have a concern about the study or I wish to complain?</u>
If you have a concern about any aspect of this study, please speak to [redacted] or their supervisor [redacted] and we will do our best to answer your query. We will acknowledge your concern within 10 working days and give you an indication of how it will be dealt with. If you remain unhappy or wish to make a formal complaint, please contact the Chair of the Research Ethics Committee who will seek to resolve the matter as soon as possible:
School of Geography and the Environment Departmental Research Ethics Committee Chair: Prof Jim Hall, Contactable via Gillian Willis, School of Geography and the Environment, University of Oxford, South Parks Road, Oxford, OX1 3QY, UK.

* 1. Please note that you may only participate in this survey if you are 18 years of age or over. If you are not, please exit the survey.

◯ Yes - I certify that I am 18 years of age or over

* 2. If you have read the information above and agree to participate with the understanding that the data (including any personal data) you submit will be processed accordingly, please check the relevant box below to get started. If not, please exit the survey.

◯ Yes - I agree to take part

### Great Crested Newt Habitat Suitability Index Users Survey

#### Part 1 of 3: Your use of the Habitat Suitability Index

1. For approximately how many years have you used the HSI for GCNs?

0 21

2. Approximately how many times per year do you use the HSI for GCNs (excluding years when not doing any)?

- 1-10
- 11-50
- >50

3. How accurate do you think the HSI is as a tool to estimate likelihood of GCN presence/absence?

Very inaccurate | Quite inaccurate | Quite accurate | Very accurate

4. How accurate do you think the HSI is as a tool to estimate abundance of GCNs?

Very inaccurate | Quite inaccurate | Quite accurate | Very accurate

### Great Crested Newt Habitat Suitability Index Users Survey

#### Part 2 of 3: Factors in the HSI

In the following questions, you will be asked to rate the accuracy of the 10 sub-indices within the HSI.

Responses of "inaccurate" will be used to infer that those sub-indices may not be suitable in their current form for inclusion in the HSI and may need changing. Response of "accurate" will infer support for the inclusion of those sub-indices in their current state. The scale from "quite" to "very" will be used to infer the strength of those views. Answers to these questions will be used in combination with ecological data to investigate ways to improve the HSI.

When answering the following questions, you may find it helpful to refer to an HSI guidance document, such as the document available here: http://www.narrs.org.uk/documents/HSI%20guidance.pdf

The first factor in the HSI is geographic location (SI1). Sites are scored according to the zone in which they occur, using a figure of the UK.

##### 1. How accurate do you think records of SI1 (Geographic Location) are?

Very inaccurate — Quite inaccurate — Quite accurate — Very accurate

○ ○ ○ ○ ○ ○ ○

##### 2. What limits the accuracy of records of SI1? (Optional)

##### 3. How accurate do you think records of SI2 (Pond Area) are?

Very inaccurate — Quite inaccurate — Quite accurate — Very accurate

○ ○ ○ ○ ○ ○ ○

The second factor is pond area (SI2). The index value is read off a graph.

##### 4. What limits the accuracy of records of SI2? (Optional)

The third factor is pond permanence (SI3). The index value is given by one of four categories (never, rarely, sometimes or annually dries out)

5. How accurate do you think records of SI3 (Pond Permanence) are?

| Very inaccurate | | Quite inaccurate | | Quite accurate | | Very accurate |
|---|---|---|---|---|---|---|
| ○ | ○ | ○ | ○ | ○ | ○ | ○ |

6. What limits the accuracy of records of SI3? (Optional)

The fourth factor is water quality (SI4). The index value is given by one of four categories (good, moderate, poor or bad).

7. How accurate do you think records of SI4 (Water Quality) are?

| Very inaccurate | | Quite inaccurate | | Quite accurate | | Very accurate |
|---|---|---|---|---|---|---|
| ○ | ○ | ○ | ○ | ○ | ○ | ○ |

8. What limits the accuracy of records of SI4? (Optional)

The fifth factor is shade (SI5). The index value is read off a graph, based on the percentage of the pond perimeter which is shaded.

9. How accurate do you think records of SI5 (Shade) are?

| Very inaccurate | | Quite inaccurate | | Quite accurate | | Very accurate |
|---|---|---|---|---|---|---|
| ○ | ○ | ○ | ○ | ○ | ○ | ○ |

10. What limits the accuracy of records of SI5? (Optional)

The sixth factor is the impact of waterfowl (SI6). The index value is given by one of three categories (absent, minor, major).

11. How accurate do you think records of SI6 (Waterfowl) are?

| Very inaccurate | | Quite inaccurate | | Quite accurate | | Very accurate |
|---|---|---|---|---|---|---|
| ○ | ○ | ○ | ○ | ○ | ○ | ○ |

12. What limits the accuracy of records of SI6? (Optional)

The seventh factor is the impact of fish (SI7). The index value is given by one of four categories (absent, possible, minor, major).

#### 13. How accurate do you think records of SI7 (Fish) are?

| Very inaccurate | | Quite inaccurate | | Quite accurate | | Very accurate |
|---|---|---|---|---|---|---|
| ○ | ○ | ○ | ○ | ○ | ○ | ○ |

#### 14. What limits the accuracy of records of SI7? (Optional)

The eighth factor is pond count (SI8). The index value is read off a graph, based on the number of ponds within 1km.

#### 15. How accurate do you think records of SI8 (Pond Count) are?

| Very inaccurate | | Quite inaccurate | | Quite accurate | | Very accurate |
|---|---|---|---|---|---|---|
| ○ | ○ | ○ | ○ | ○ | ○ | ○ |

#### 16. What limits the accuracy of records of SI8? (Optional)

The ninth factor is terrestrial habitat (SI9). The index value is given by one of four categories (good, moderate, poor, none). In the original version of the HSI, the index value is read off a graph, based in the amount of good terrestrial habitat within 500m - if you use this method please indicate this in the answer to question 18.

#### 17. How accurate do you think records of SI9 (Terrestrial Habitat) are?

| Very inaccurate | | Quite inaccurate | | Quite accurate | | Very accurate |
|---|---|---|---|---|---|---|
| ○ | ○ | ○ | ○ | ○ | ○ | ○ |

#### 18. What limits the accuracy of records of SI9? (Optional)

The tenth factor is macrophyte cover (SI10). The index value is read off a graph.

#### 19. How accurate do you think records of SI10 (Macrophyte Cover) are?

| Very inaccurate | | Quite inaccurate | | Quite accurate | | Very accurate |
|---|---|---|---|---|---|---|
| ○ | ○ | ○ | ○ | ○ | ○ | ○ |

20. What limits the accuracy of records of SI10? (Optional)

UNIVERSITY OF OXFORD

#### Great Crested Newt Habitat Suitability Index Users Survey

##### Part 3 of 3: Other factors

1. Do you think there are other environmental factors that may impact the suitability of ponds for GCNs that are not currently included in the HSI? If so, please list below.

2. Do you have other suggestions for the improvement of the HSI for GCNs? If so, please list below.

3. In what capacity do you use the HSI for GCNs? (If more than one category applies, please select the capacity under which you usually use the index)

- ◯ Volunteer – independent
- ◯ Volunteer - ecological consultancy or similar
- ◯ Volunteer - conservation organisation
- ◯ Other (please specify)
- ◯ Professional - conservation organisation
- ◯ Professional - ecological consultancy or similar

**Supp.Mat.3: Additional statistical information from post-hoc tests**

Supp.Mat.3a: Statistical information from posthoc test looking at relationship between HSI categories and ponds with GCN presence/absence.

| | | HSI Categories | | | | |
|---|---|---|---|---|---|---|
| | | Poor | Below Average | Average | Good | Excellent |
| **Residual** | **Absent** | 11.914 | 5.532 | -2.2618 | -7.891 | -7.226 |
| | **Present** | -11.914 | -5.532 | 2.2618 | 7.891 | 7.226 |
| **P-value** | | <0.001 | <0.001 | 0.237 | <0.001 | <0.001 |

Supp.Mat.3b: Statistical information from posthoc test looking at relationship between HSI categories, based on arithmetic HSI and new category division, and GCN presence/absence.

| | | HSI Categories | | | | |
|---|---|---|---|---|---|---|
| | | Poor | Below Average | Average | Good | Excellent |
| **Residual** | **Absent** | 14.007 | 2.889 | -0.696 | -6.374 | -9.0779 |
| | **Present** | -14.007 | -2.889 | 0.696 | 6.374 | 9.0779 |
| **P-value** | | <0.001 | 0.039 | 1.000 | <0.001 | <0.001 |

Supp.Mat.3c: Statistical information from posthoc test looking at relationship between HSI categories, with HSI scores recalculated with SI4 (Water quality) excluded, and GCN presence/absence.

| | | HSI Categories with SI4 excluded | | | | |
|---|---|---|---|---|---|---|
| | | Poor | Below Average | Average | Good | Excellent |
| **Residual** | **Absent** | 12.279 | 4.227 | -1.082 | -6.182 | -10.604 |
| | **Present** | -12.279 | -4.227 | 1.082 | 6.182 | 10.604 |
| **P-value** | | <0.001 | <0.001 | 1.000 | <0.001 | <0.001 |

Supp.Mat.3d: Statistical information from posthoc test looking at relationship between HSI categories, with HSI scores recalculated with SI6 (Waterfowl) excluded, and GCN presence/absence.

| | | **HSI Categories with SI6 excluded** | | | | |
|---|---|---|---|---|---|---|
| | | **Poor** | **Below Average** | **Average** | **Good** | **Excellent** |
| **Residual** | **Absent** | 15.379 | 5.031 | -4.767 | -8.395 | -8.105 |
| | **Present** | -15.379 | -5.031 | 4.767 | 8.395 | 8.105 |
| **P-value** | | <0.001 | <0.001 | <0.001 | <0.001 | <0.001 |

Supp.Mat.3e: Statistical information from posthoc test looking at relationship between HSI categories, with HSI scores recalculated with SI4 (Water quality) and SI6 (Waterfowl) excluded, and GCN presence/absence.

| | | **HSI Categories with SI4 and SI6 excluded** | | | | |
|---|---|---|---|---|---|---|
| | | **Poor** | **Below Average** | **Average** | **Good** | **Excellent** |
| **Residual** | **Absent** | 16.173 | 5.589 | -4.253 | -7.191 | -9.719 |
| | **Present** | -16.173 | -5.589 | 4.253 | 7.191 | 9.719 |
| **P-value** | | <0.001 | <0.001 | <0.001 | <0.001 | <0.001 |

Supp.Mat.3f: Statistical information from posthoc test looking at relationship between HSI categories, with HSI scores recalculated with new SI2 (Pond size) scores, and GCN presence/absence.

| | | **HSI categories** | | | | |
|---|---|---|---|---|---|---|
| | | **Poor** | **Below Average** | **Average** | **Good** | **Excellent** |
| **Residual** | **Absent** | 11.297 | 7.403 | 4.545 | -8.089 | -10.041 |
| | **Present** | -11.297 | -7.403 | -4.545 | 8.089 | 10.041 |
| **P-value** | | <0.001 | <0.001 | <0.001 | <0.001 | <0.001 |

Supp. Mat. 3g: Statistical information from posthoc test looking at relationship between HSI category with HSI scores recalculated using newly modified HSI and GCN presence/absence.

| | | New HSI with combined modifications | | | | |
|---|---|---|---|---|---|---|
| | | **Poor** | **Below Average** | **Average** | **Good** | **Excellent** |
| **Residual** | **Absent** | 20.525 | 4.753 | -2.011 | -10.791 | -9.718 |
| | **Present** | -20.525 | -4.753 | 2.011 | 10.791 | 9.718 |
| **P-value** | | p<0.001 | p<0.001 | 0.443 | p<0.001 | p<0.001 |

Supp. Mat. 3h: Statistical information from binomial generalised linear model, looking at HSI scores calculated with both the newly modified HSI and existing HSI, and GCN presence/absence, on the independent data-set for cross validation and the main dataset.

| | **Original dataset - old HSI** | **Original dataset - new HSI** | **Verification dataset - old HSI** | **Verification dataset - new HSI** |
|---|---|---|---|---|
| **Effect size** | 0.736 | 5.266 | 0.7759 | 1.411 |
| **Direction** | Positive | Positive | Positive | Positive |
| **P value** | <0.001 | <0.001 | 0.169 | 0.156 |
| **R2 (calculated from glm as 1-(residual deviance/null deviance)** | 0.029 | 0.038 | 0.041 | 0.043 |

Supp. Mat. 3i: Statistical information from posthoc test looking at relationship between HSI categories, with HSI scores calculated with both the newly modified HSI and existing HSI, and GCN presence/absence, on the independent data-set for cross validation.

| | | | HSI Categories | | | | |
|---|---|---|---|---|---|---|---|
| | | | **Poor** | **Below Average** | **Average** | **Good** | **Excellent** |
| **Original data - old HSI** | **Residual** | **Absent** | 11.914 | 5.532 | -2.262 | -7.891 | -7.226 |
| | | **Present** | -11.914 | -5.532 | 2.262 | 7.891 | 7.226 |
| | **P value** | | <0.001 | <0.001 | 0.237 | <0.001 | <0.001 |
| **Original data - new HSI** | **Residual** | **Absent** | 20.525 | 4.753 | -2.011 | -10.791 | -9.718 |
| | | **Present** | -20.525 | -4.753 | 2.011 | 10.791 | 9.718 |
| | **P value** | | <0.001 | <0.001 | 0.443 | <0.001 | <0.001 |
| **Verification data - old HSI** | **Residual** | **Absent** | 1.221 | 0.543 | 0.819 | -1.353 | -1.258 |
| | | **Present** | -1.221 | -0.543 | -0.819 | 1.353 | 1.258 |
| | **P value** | | 1 | 1 | 1 | 1 | 1 |
| **Verification data - new HSI** | **Residual** | **Absent** | 1.074 | 0.883 | -0.079 | -0.543 | -2.124 |
| | | **Present** | -1.074 | -0.883 | 0.079 | 0.543 | 2.124 |
| | **P value** | | 1 | 1 | 1 | 1 | 0.335 |

**Supp.Mat.4: Qualitative survey results**

Hierarchy maps with quantification of themes and additional detailed findings.

Supp.Mat.4a: SI1-10

Hierarchy map of nodes created from textual analysis of survey answers to the question "What limits the accuracy of SI(x) records?" Numbers indicate the number of references to that specific concept. The number of answers to the question is given in the top node.

SI1:

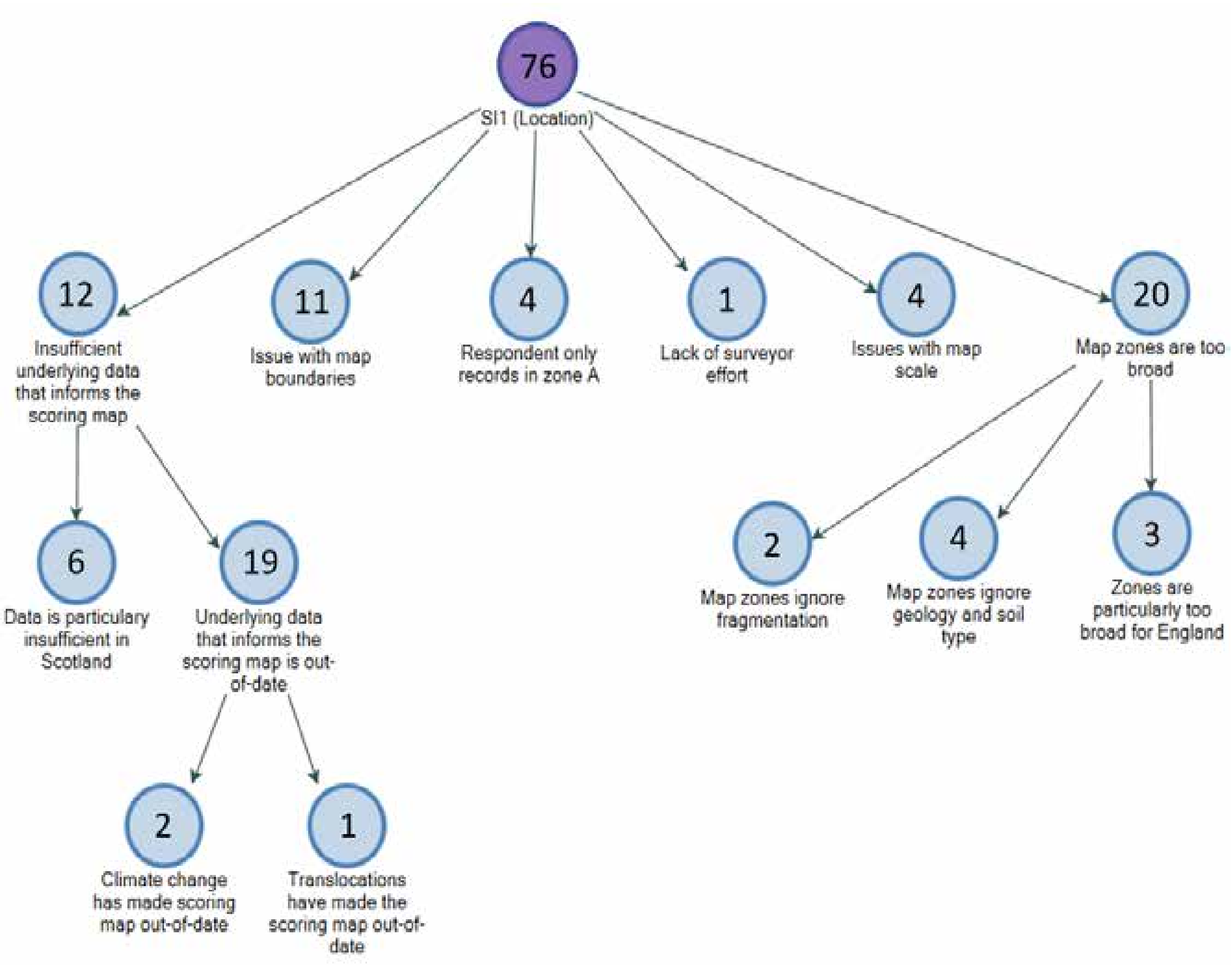


Of the comments on limitations of SI1 accuracy, most responses related to the zones being too broad or the underlying data being insufficient. Some comments suggested that the data was out-of-date, especially for Scotland. Several respondents suggested that the map image was hard to read, particularly for identifying the boundaries.

SI2:

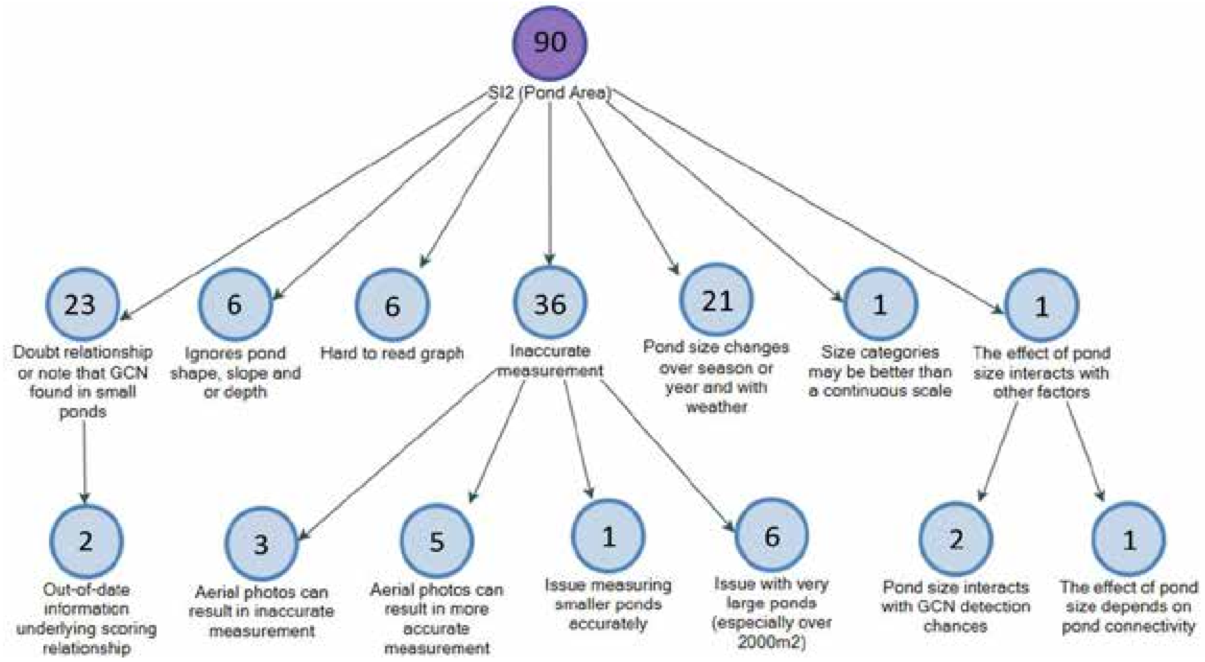


The main criticism of SI2 accuracy related to imprecise measurement, with respondents often criticising or supporting certain methods. Many respondents indicated that records were often estimated rather than measured. Numerous respondents gave examples of finding GCNs in small waterbodies. Several respondents noted that the graph only covers waterbodies smaller than 2000m2. Some suggested consideration of depth or other pond shape variables.

SI3:

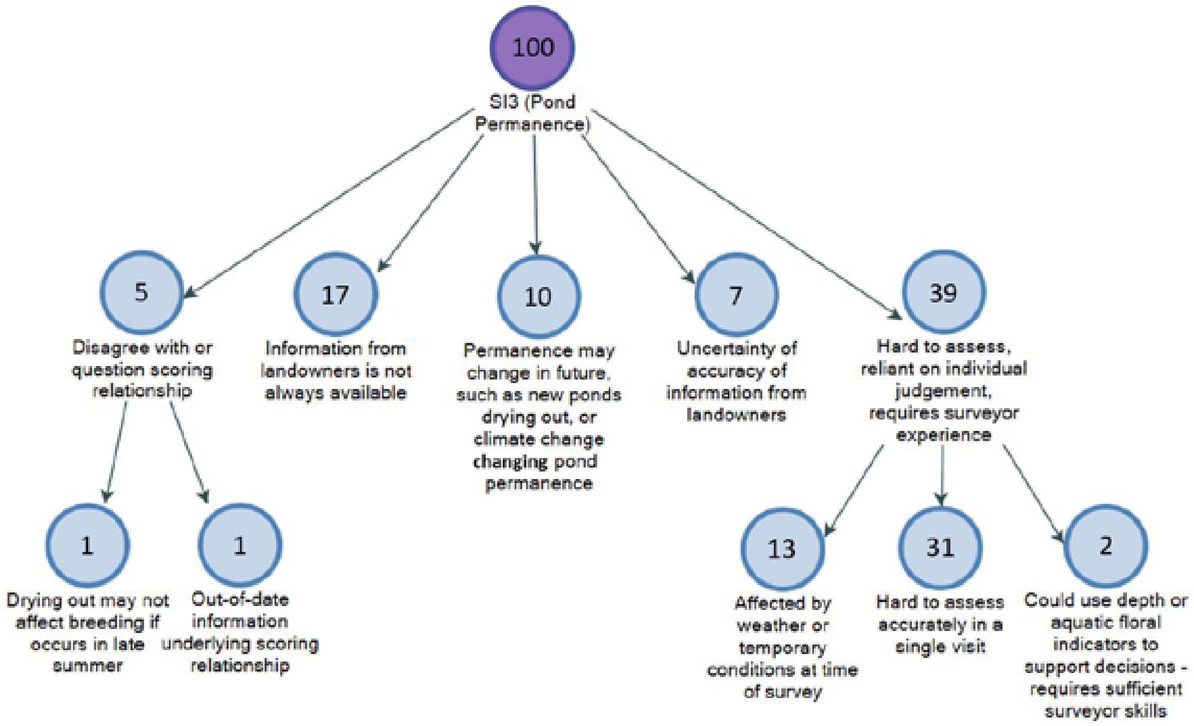


Most of the responses around limitations to SI3 accuracy related to the difficulty and subjectivity of assessing pond permanence. Information from local people may be relied upon but this may be inaccurate.

SI4:

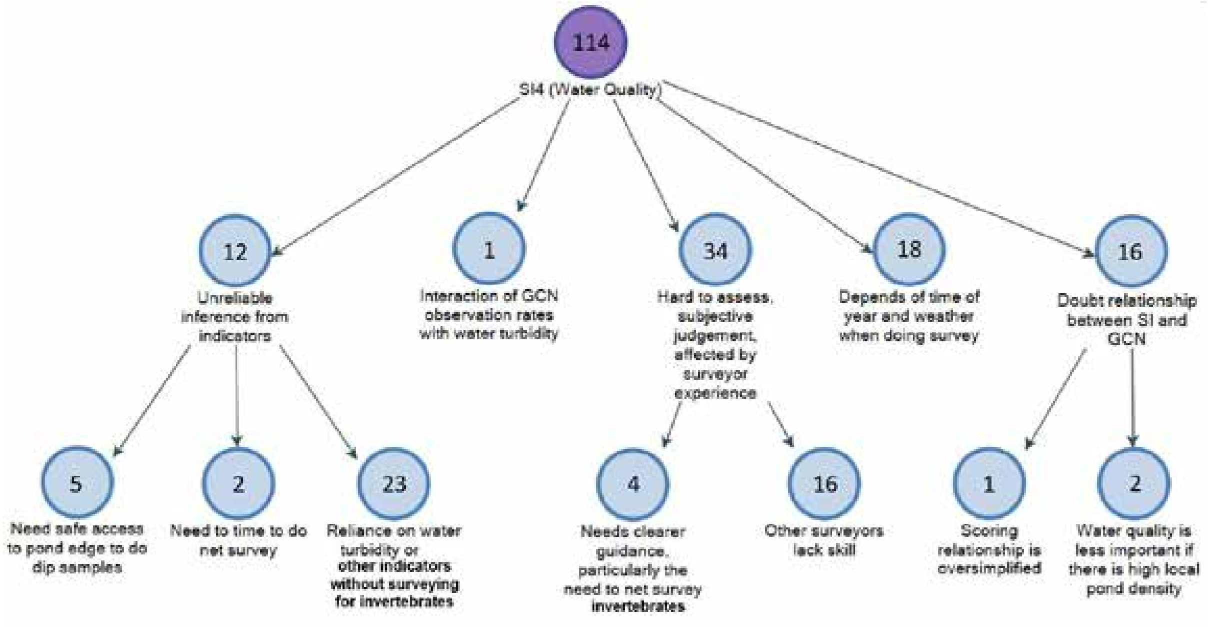


Many responses on SI4 accuracy related to the subjectivity, and suggested records further vary between surveyors due to experience and skill. Many noted the difficulty of assessing water quality from surface visual assessments and suggested a need to do a net survey of invertebrates. Access and time were listed as reasons for not doing a dip survey. There were many references to the time of year impacting records.

SI5:

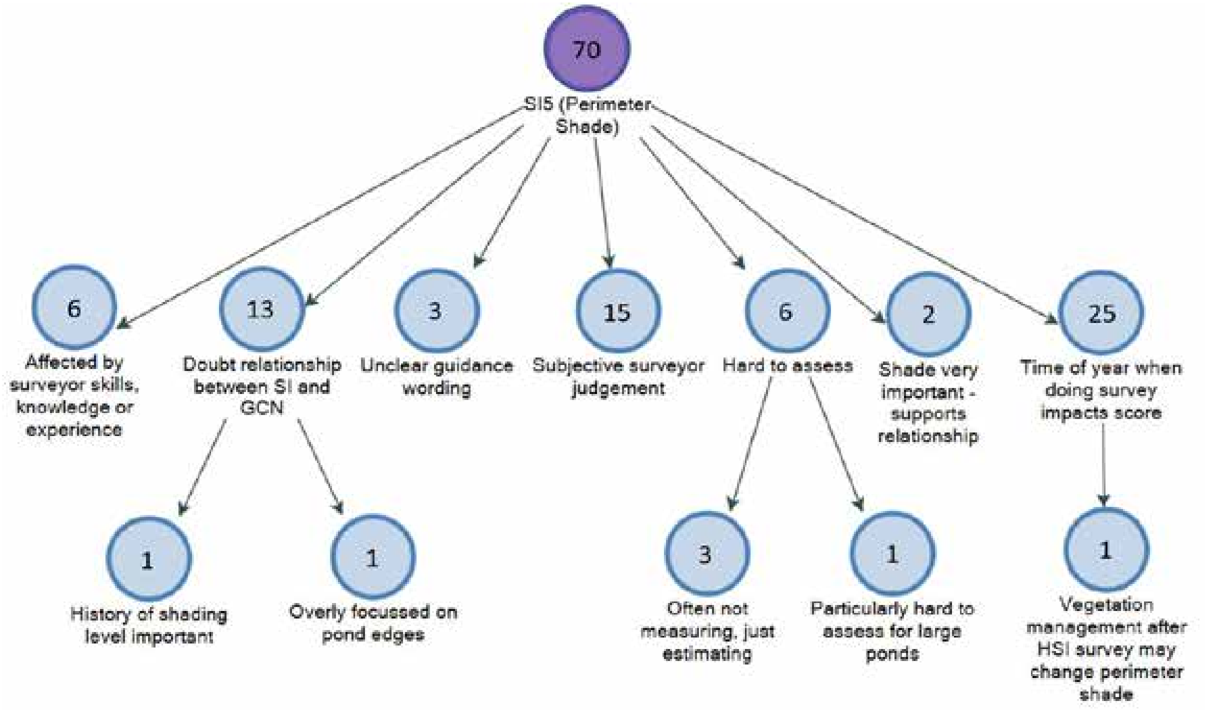


The most common comment regarding SI5 accuracy limitations was that shade varied across time, so a single site visit may not give an accurate answer, especially if surveying in winter. Another common response was to note the subjectivity of assessment, or the variability in records based on surveyor skills. Several responses suggested difficulties in measurement, with records often estimated.

SI6:

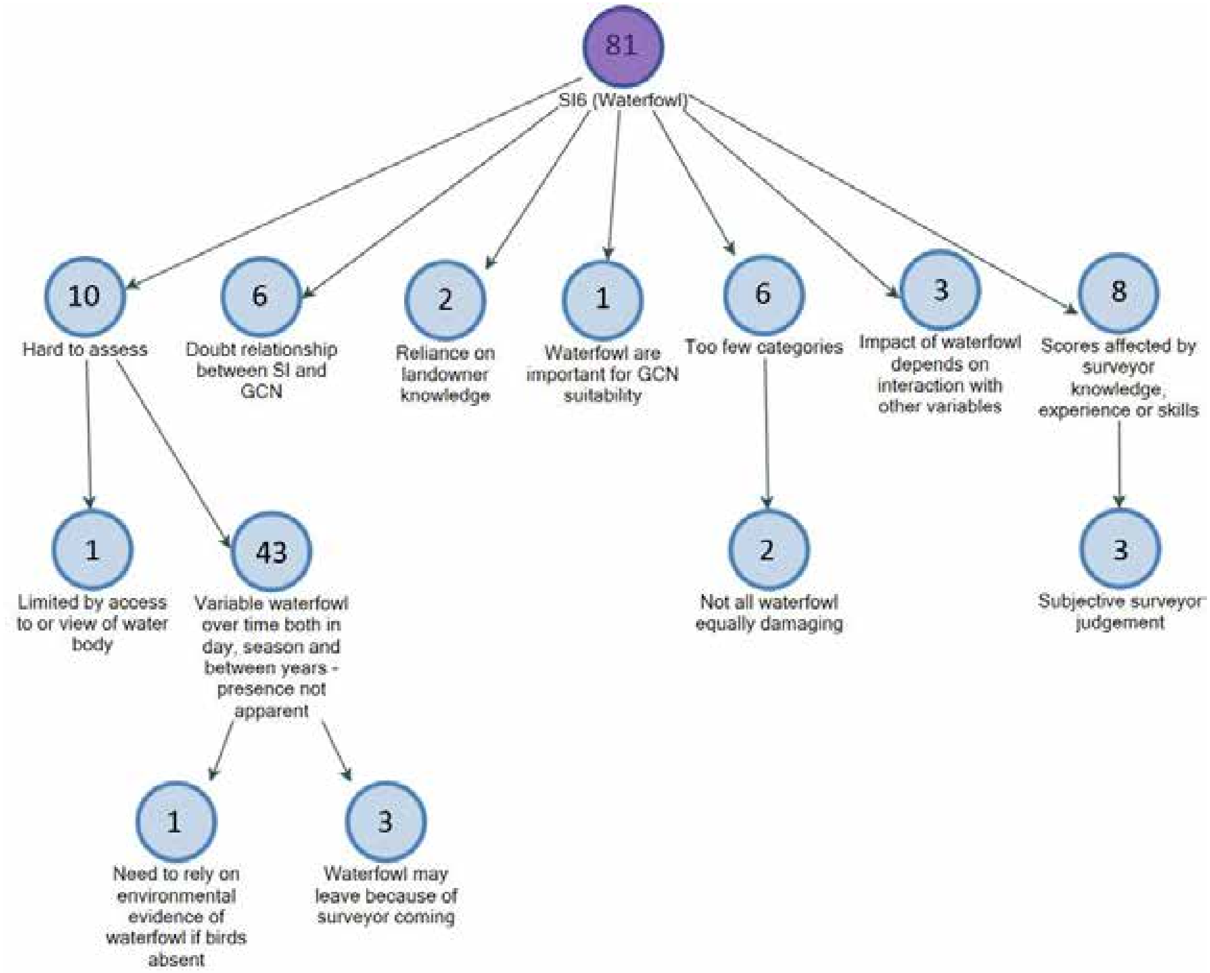


Most responses to the question on SI6 mentioned the temporal variability of waterfowl presence, noting changes within a day, across the season, and between years. Locals may be consulted, but they may not have reliable information. Several people noted the reliance on surveyor skill and subjective judgement. Several respondents suggested having an intermediate category, and having greater guidance on species to include.

SI7:

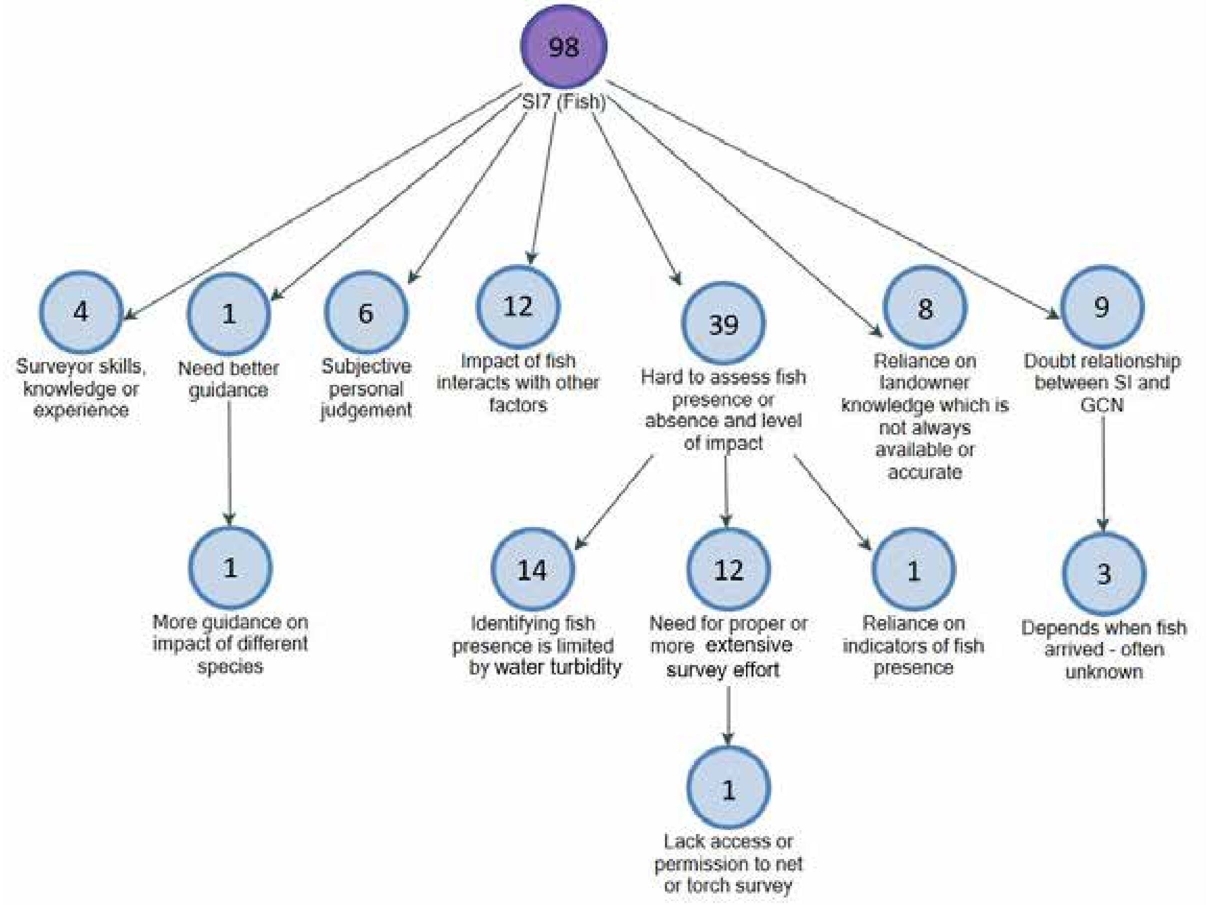


Many respondents gave a generic comment about the difficulty of determining fish presence for SI7, with some associating this with water turbidity. Several criticised relying on landowners, while others criticised surveyors' skills. Several comments indicated subjective judgement.

SI8:

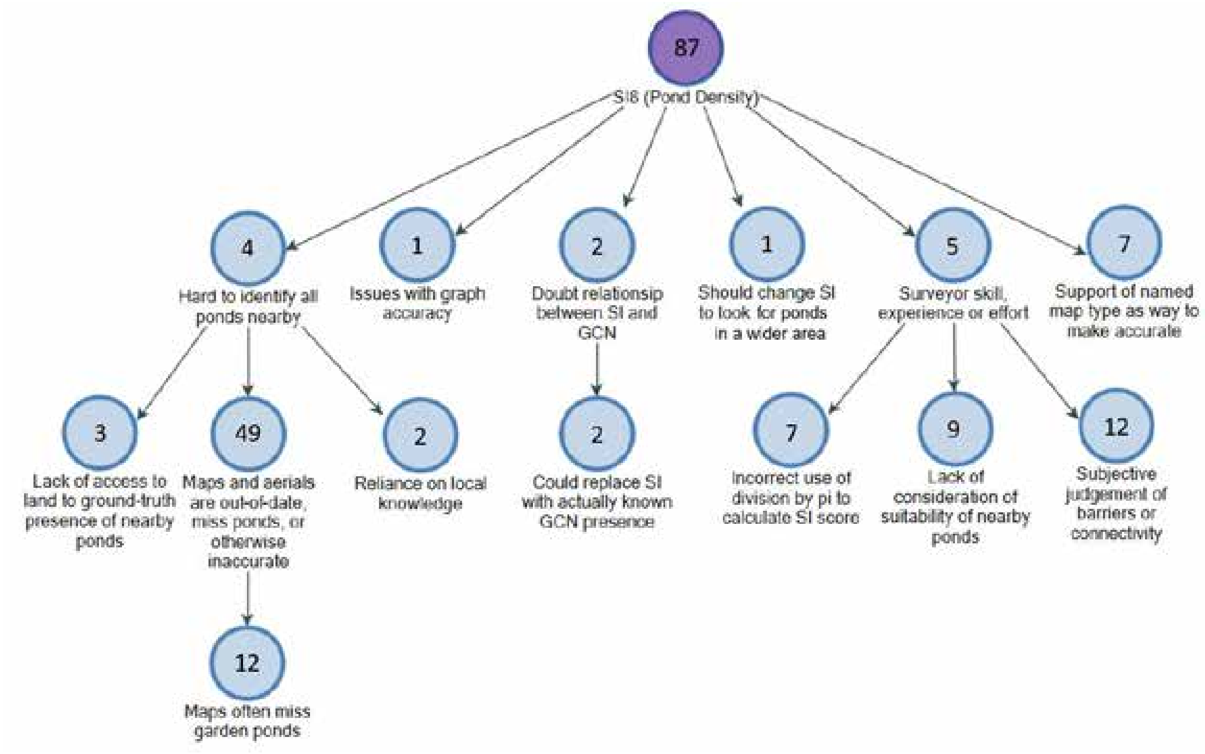


A very common issue identified with SI8 records was that maps and aerial photography used to identify local ponds may be inaccurate, particularly for garden ponds. Some respondents noted the importance of checking the nearby ponds for suitability and connectivity, which was thought to be rather subjective. There were a few mentions of surveyor skill, with seven respondents noting that other surveyors sometimes forget to divide by pi before reading the SI value off the graph. A useful suggestion was to create a recalibrated graph to remove the need for division.

SI9:

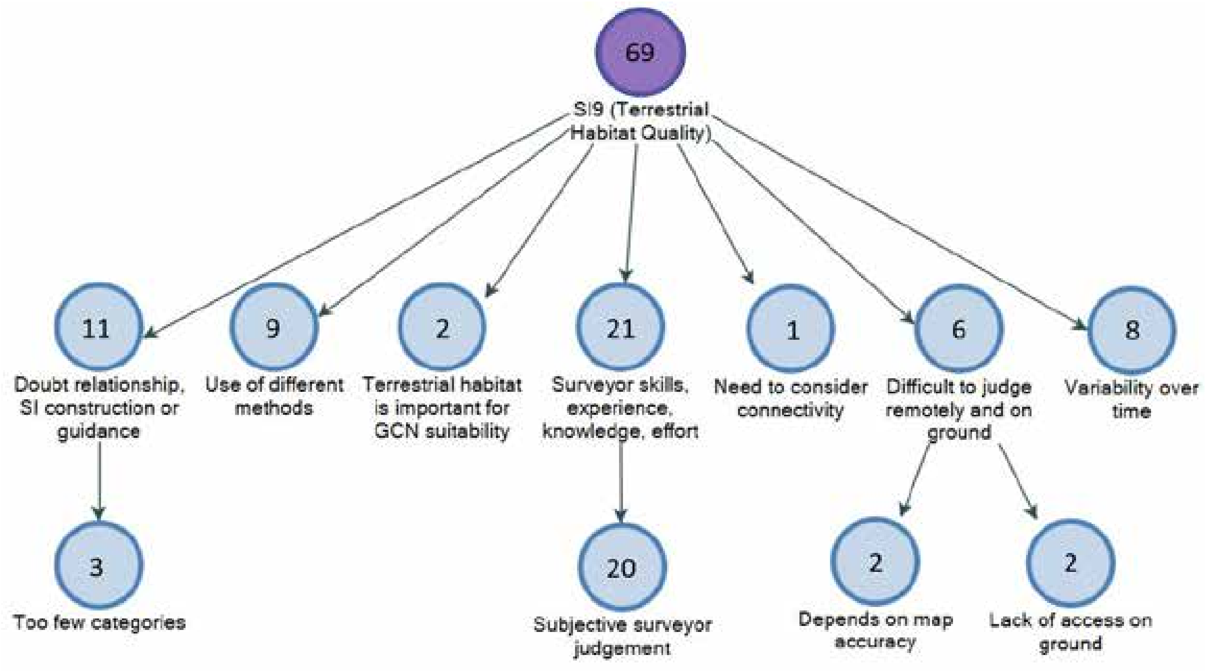


For SI9 accuracy, most respondents' answers focussed on subjectivity of records, impacted by surveyor skills, experience, knowledge or effort. Several said the categories were too broad. Other difficulties in assessment came from lack of physical access or limitations of maps. There were several comments on the temporal variability of habitat quality.

SI10:

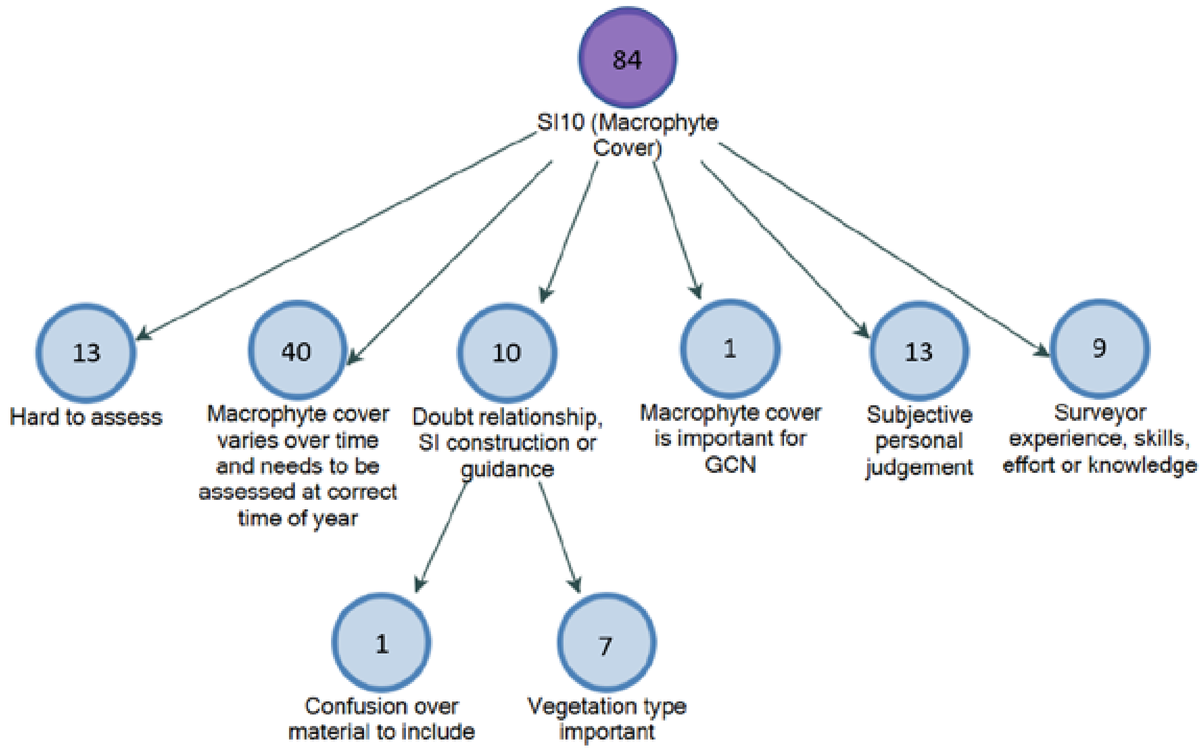


Many respondents suggested practical limitations to SI10 record accuracy. Almost half the responses noted the temporal variation of macrophyte cover. Others noted that they often have to conduct HSI surveys year-round, with some suggesting this was acceptable and others not. Another common response focussed on subjectivity, with many commenting on the impact of skill. Several respondents suggested a need for more detailed guidance specifying which macrophytes to include. Many comments suggested macrophyte cover was not a useful measure, and instead egg-laying material would be more appropriate.

Supp.Mat.4b: Other Factors:

Hierarchy map of nodes created from textual analysis of survey answers on additional factors to include in the HSI. Categories added after coding are shaded green to aid visualisation. Numbers indicate the number of references to that specific concept.The number of answers to the question is given in the purple central node.

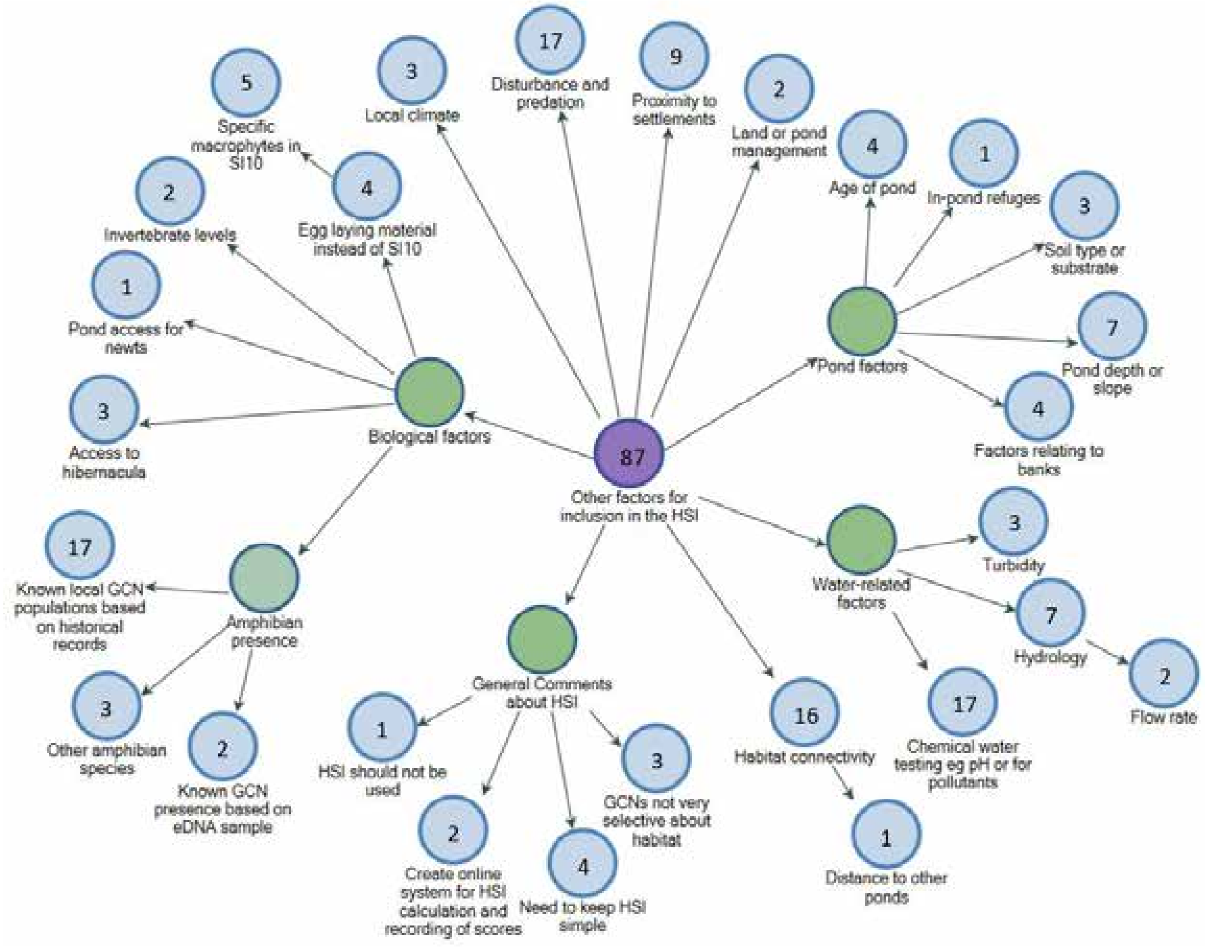


Of the 87 responses regarding possible additional factors, the most frequent suggestions (with 17 comments each) were: known GCN populations in the area; chemical water tests; and disturbance or predation. Comments on disturbance predominantly referenced dogs and people, with occasional mentions of other pets and non-native species. These comments relate to mentions of proximity to settlements, which usually were made regarding disturbance, risk from roads or general habitat unsuitability. Comments around water chemistry suggested testing for pH, fertilisers, minerals, salinity or pollutants.

Although habitat connectivity should be considered under SI8 and SI9, there seemed a desire to address this more directly. Several suggestions concerned the pond shape, age, and substrate, with pond depth suggested by several respondents. A few comments were made relating to water turbidity and hydrology. A number of comments relating directly to pond biology were made, such as presence of other amphibians, invertebrates or specific macrophytes. Few people discussed the practical complexities including these additional variables would bring. Four people noted the need to keep the HSI simple to use.

Supp.Mat.4c: Suggestions for improvement:

Hierarchy map of nodes created from textual analysis of survey answers on suggestions for HSI improvement. Categories added after coding are shaded green to aid visualisation. Numbers indicate the number of references to that specific concept. The number of answers to the question is given in the purple central node.

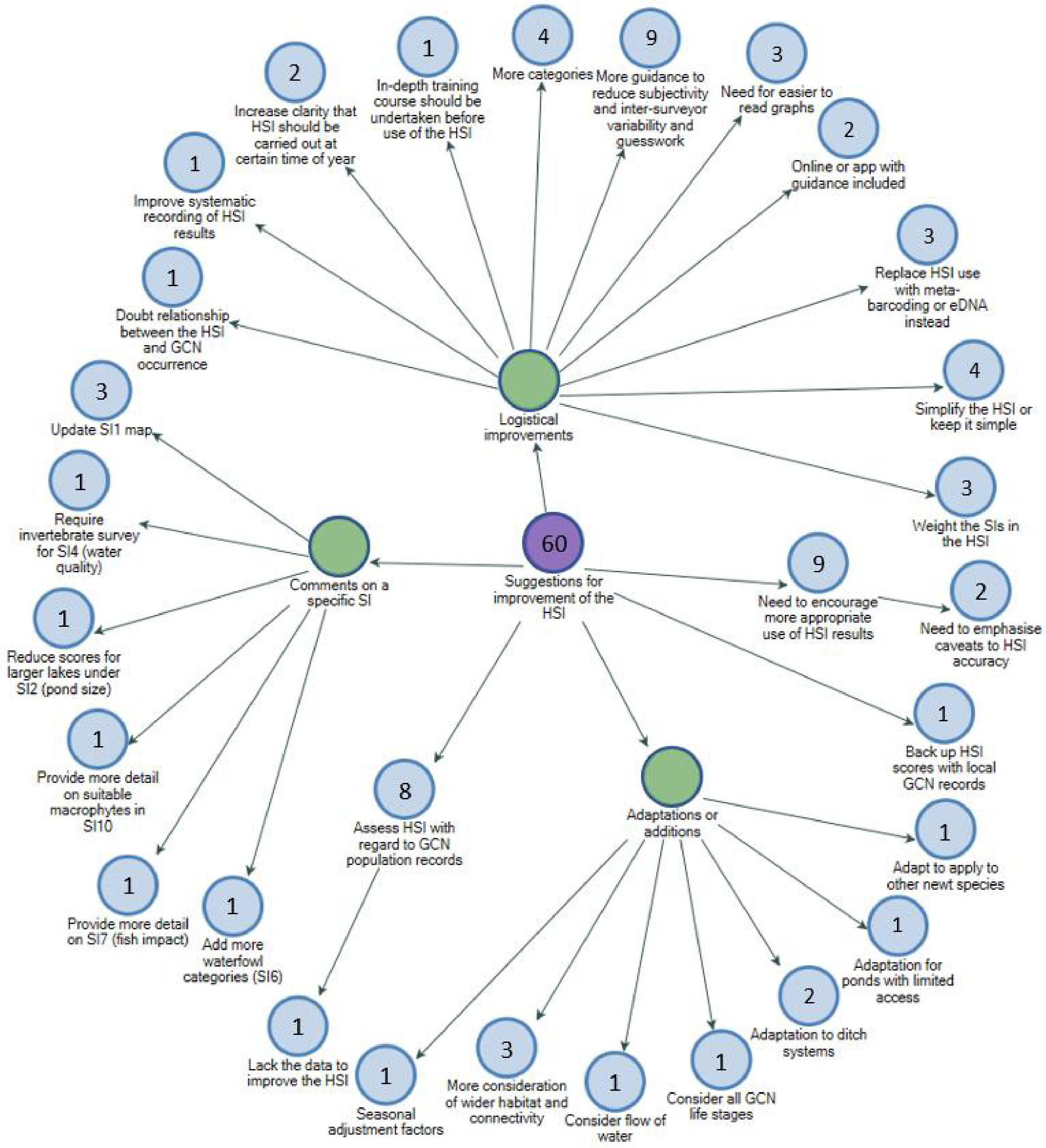


Of the 60 responses to this question, a common response was that the HSI needs to be used in a more appropriate manner (not to substitute for GCN surveys), and the guidance could emphasise this. Another common suggestion was for better guidance to reduce subjectivity.

A frequent suggestion was to incorporate local GCN records or replace the HSI with eDNA surveys. Many suggestions were made by only one or a few respondents, with much variation in the suggestions provided. Other suggestions included adaptations or additions such as considering other newt species, water flow, ditch systems or connectivity. Several logistical suggestions were made such as improving recording, training, guidance and graphs, and developing an online system. Some comments referenced specific SIs, often reiterating comments given earlier.

**Supp.Mat.5: Line formulae for new graphs**

New SI2 graph:

Where *y*=SI2 score and *x*= ponds size.

For ponds <7m$^2$:

$$y = \frac{1}{28}x + \frac{1}{2}$$

For ponds 7-20m$^2$:

$$y = \frac{1}{52}x + \frac{8}{13}$$

For ponds 20-400m$^2$:

$$y = 1$$

For ponds 400m-3000m$^2$:

$$y = -\frac{1}{10400}x + \frac{27}{26}$$

For ponds ≥3000m$^2$:

$$y = -\frac{1}{76000}x + \frac{15}{19}$$

SI5 graph:

Where *y*=SI2 score and *x*= ponds size.

For pond shade <60%:

$$y = 1$$

For pond shade ≥60%:

$$y = -\frac{2}{100}x + \frac{22}{10}$$

New SI8 graph:

Where *y*=SI8 score and *x*= number of ponds within 1km:

For ponds with >12 ponds within 1km:

$$y = 1$$

For ponds with ≤12 ponds within 1km:

$$y = \frac{3}{40}x + \frac{1}{10}$$

SI10 graph:

Where *y*=SI10 score and *x*= percentage macrophyte cover:

For ponds with <70% macrophyte cover:

$$y = \frac{1}{100}x + \frac{3}{10}$$

For ponds with 70-80% macrophyte cover:

$$y = 1$$

For ponds with >80% macrophyte cover:

$$y = -\frac{1}{100}x + \frac{18}{10}$$

**Figure S1:** Perceived accuracy of the HSI when used in the context of reflecting GCN abundance is lower than the perceived accuracy of the HSI when used in the context of reflecting GCN presence/absence. Bar charts show frequency of respondents selecting each accuracy rating option in answer to the survey questions on the accuracy of records of the HSI when used to reflect GCN presence/absence (a) or abundance (b).

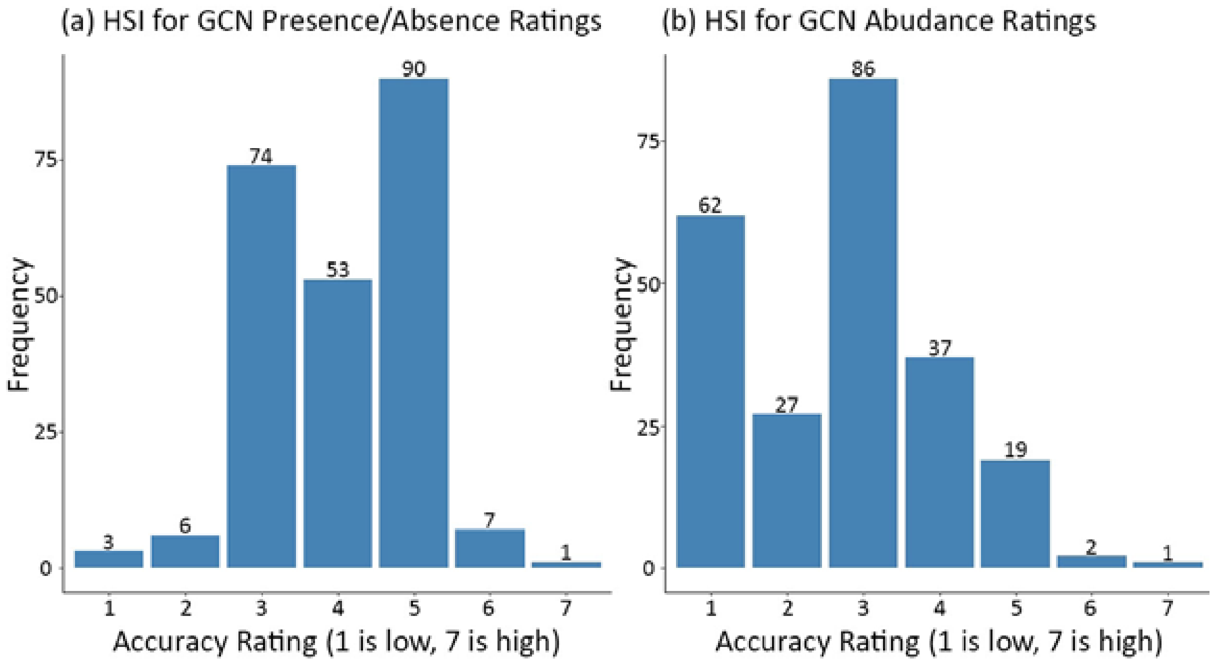

**Figure S2:** Proportion of sites with GCN presence appears to peak at intermediate values of logged pond size. Proportion of ponds with GCNs present is plotted against logged pond size (m2) and a smooth line fitted to infer the relationship between pond size and GCN presence. Some sizes with only one record have a proportion of 1 or 0, whereas sizes with multiple records can have proportions falling in between these values.

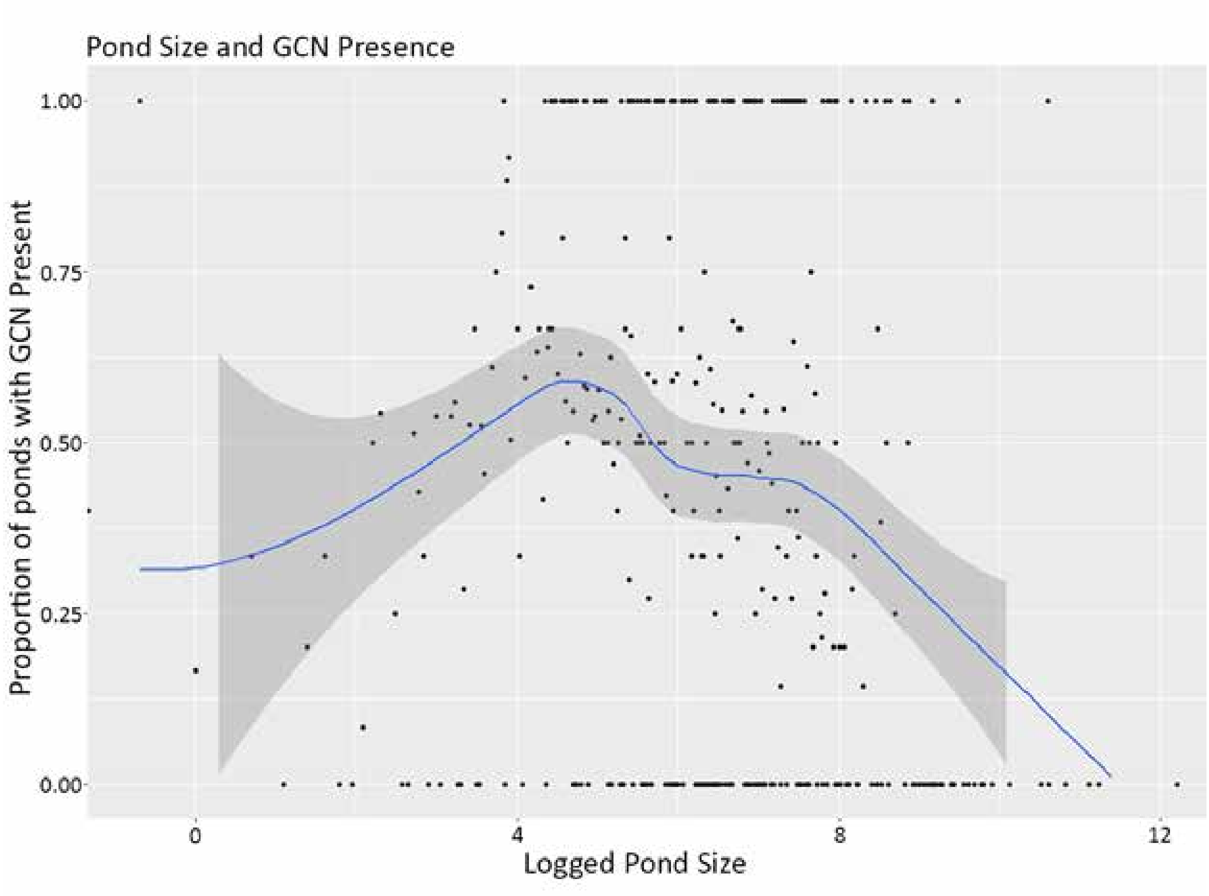

**Figure S3**: New proposed relationship for calculating SI2 scores from pond size. Note the change in scale between ponds of size ≤1000m$^2$ and pond sizes of ≥1000m$^2$, between sections (a) and (b). Ease of calculating SI2 scores from pond size is facilitated by the provision of line formulae in Supp.Mat.5.

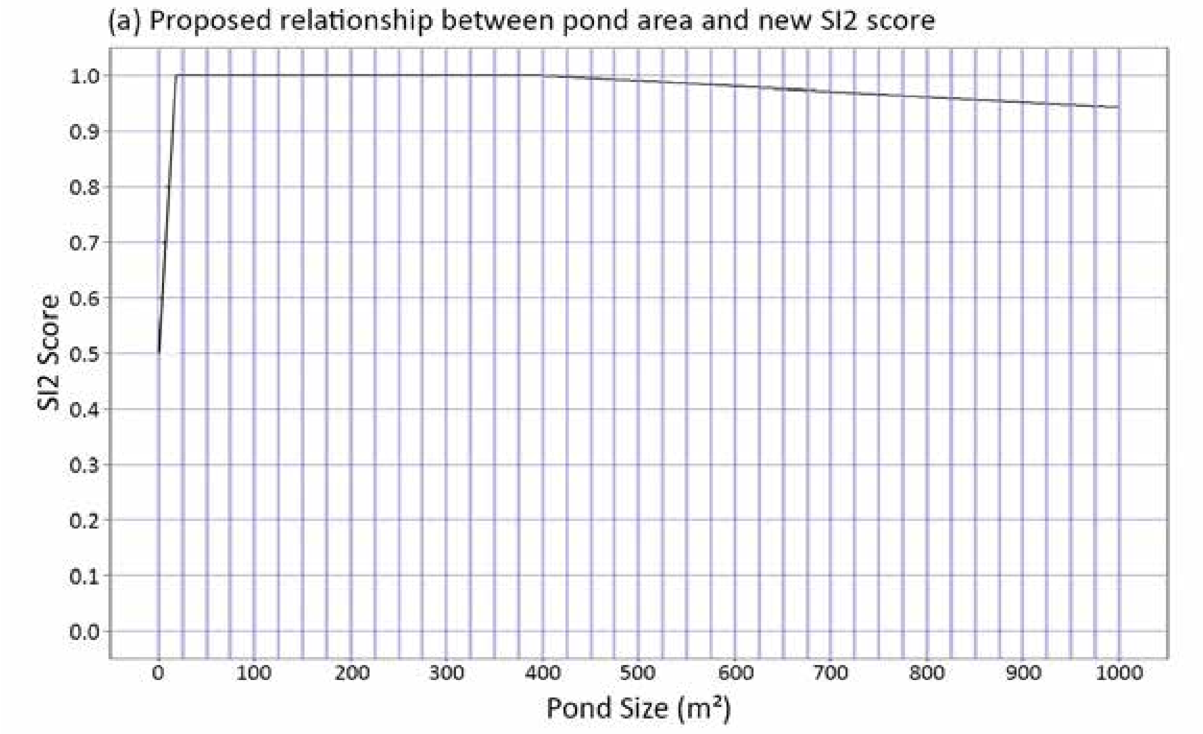


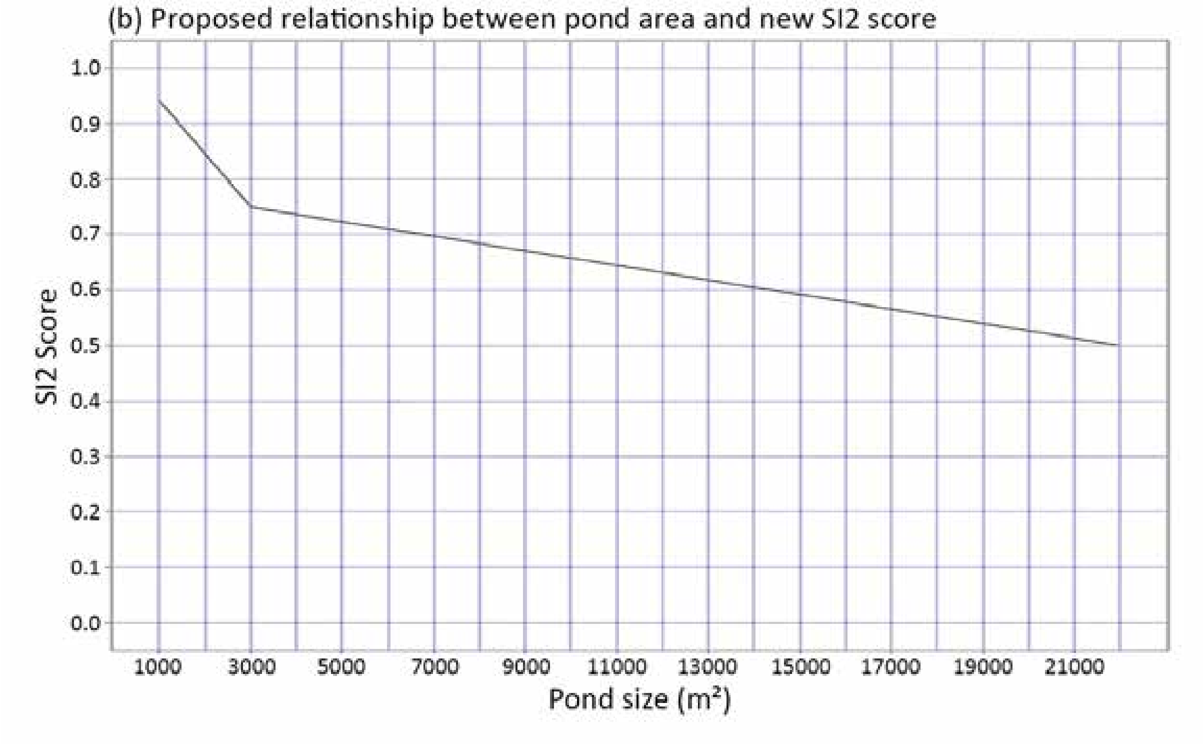

**Figure S4:** New graph to read off SI5 scores based on percentage of shoreline shade.

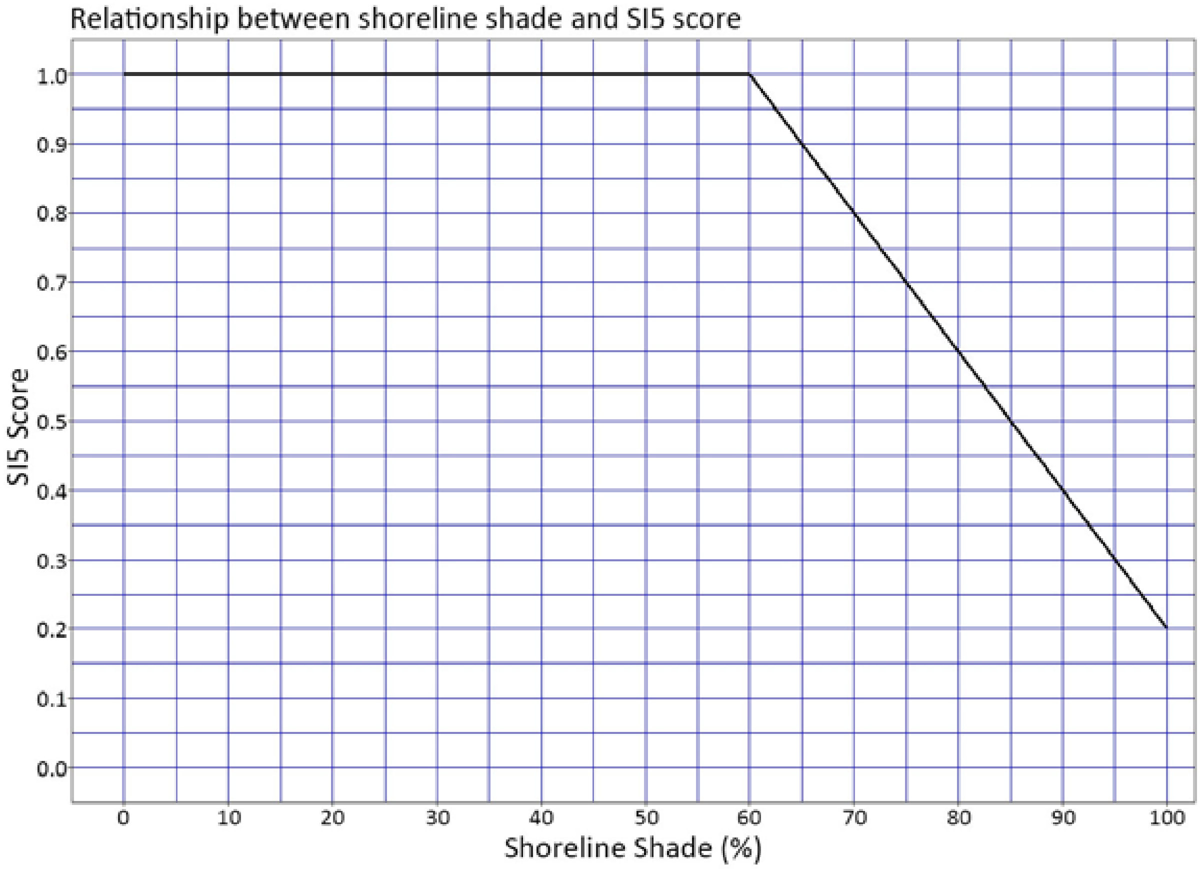

**Figure S5:** New graph to read off SI8 (Pond density) scores without the need to divide by pi first. Sites with more than 12 ponds within 1km get a score of 1. Sites with no ponds within 1km get a score of 0.1.

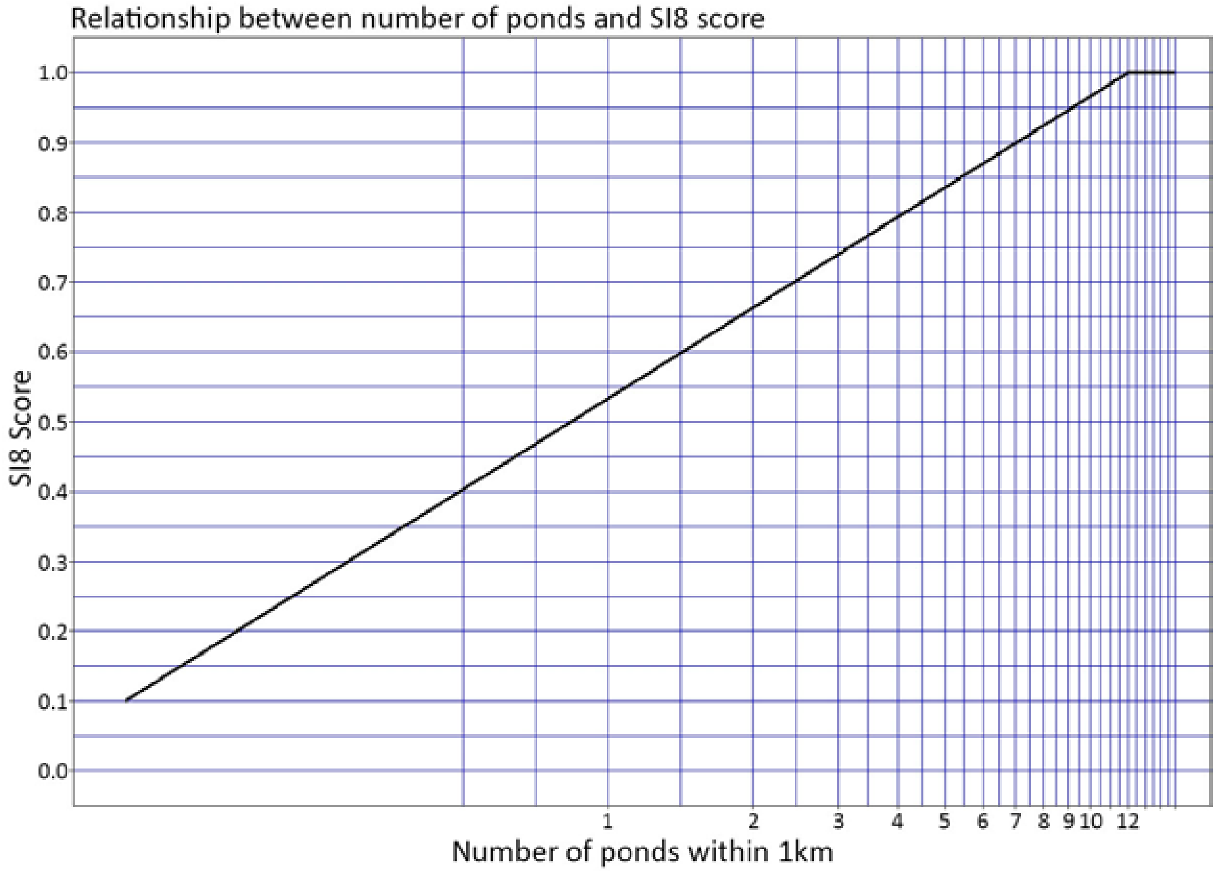

**Figure S6:** New graph to read off SI10 scores based on percentage of macrophyte cover.

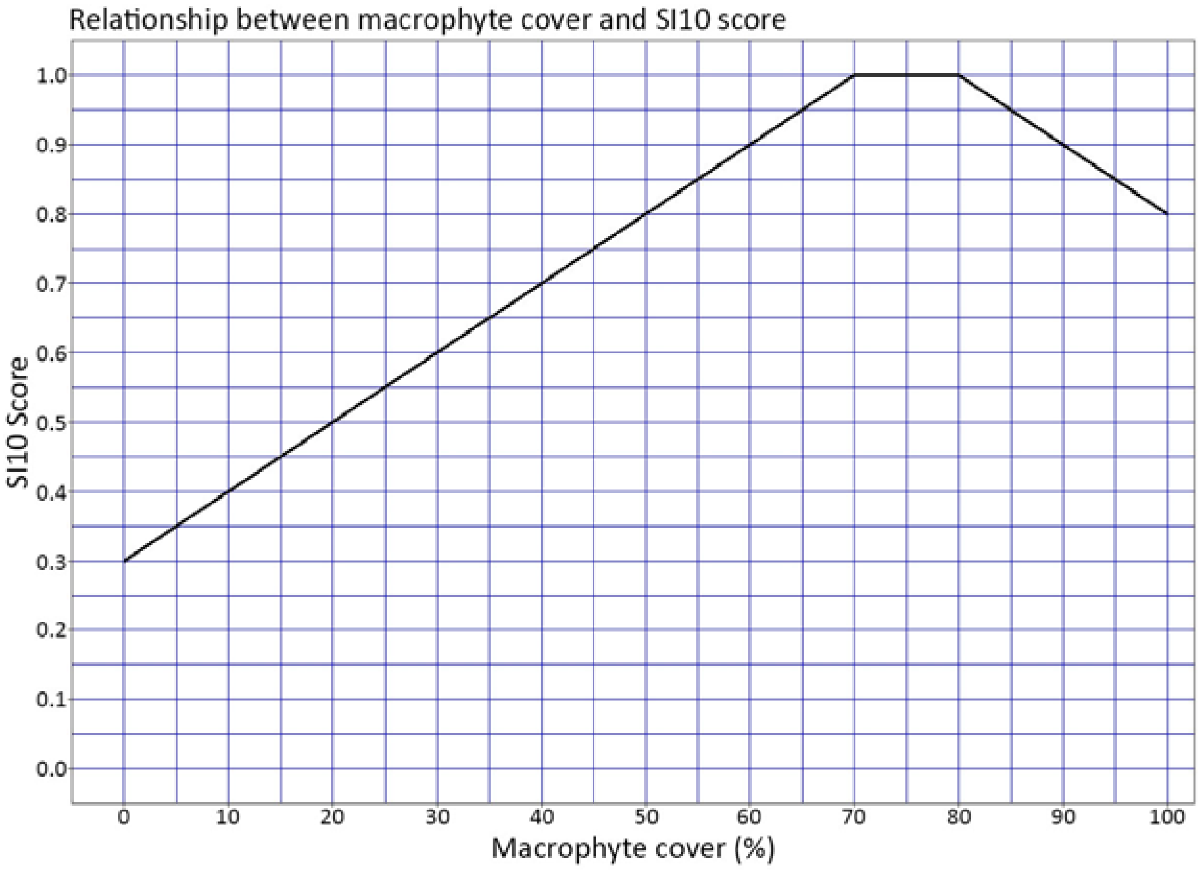
