## Supplementary material for "An improved habitat suitability index for the great crested newt": Cover Page

**Title: Improving the great crested newt habitat suitability index**

A manuscript for consideration for publication in the *Herpetological Journal* as a full-length research article.

**Contact author:**

**Author Contributions:**

Emily Seccombe: conceptualisation, methodology, formal analysis, investigation, data curation, writing (original draft preparation, review and editing), visualisation of published work, project administration.

Roberto Salguero-Gómez: Writing (review and editing), supervision.

**Ethical Statement:**

This paper abides by the British Herpetological Society: Ethical Policy and Guidelines. The research did not involve fieldwork or use of laboratory animals - all amphibian data used was pre-existing data collected by other organisations. The research involved an online survey of over 18 year olds, for which the University of Oxford's ethics policies were followed, with permission obtained from the Central University Research Ethics Committee (CUREC).
