## Supplementary material for "An improved habitat suitability index for the great crested newt": Figures

Figure 1: The perceived accuracy of suitability index (SI) records for the great crested newt (GCN) by survey respondents varies across all 10 categories (Table 1). Bar charts show the frequency of our sampled 288 respondents selecting each accuracy rating option (from 1 to 7, where 1 is the lowest) in answer to the survey questions on the accuracy of records of SIs 1-10 (a-j).

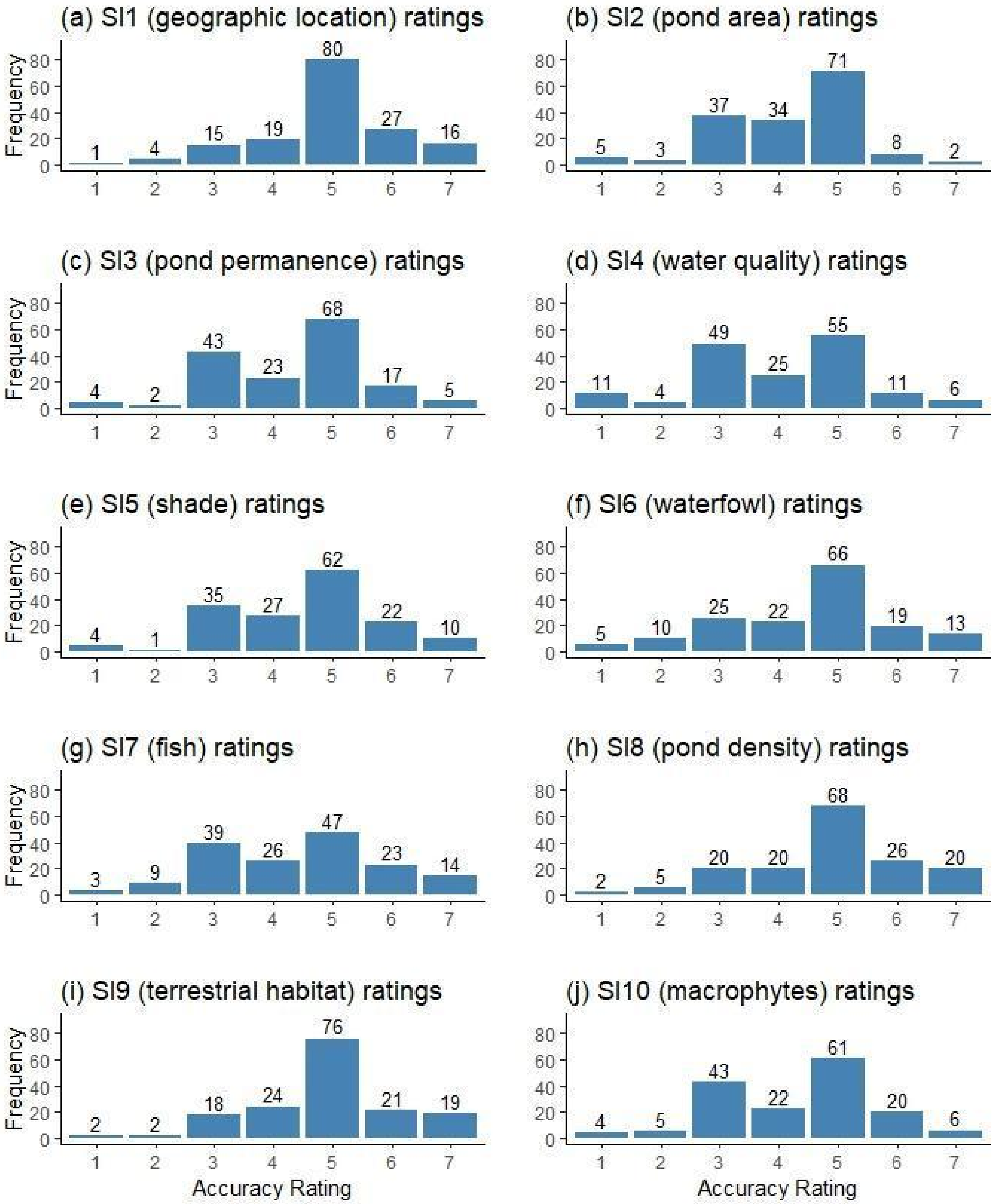

Figure 2: Minimal differences in scores of continuous suitability indexes (SIs) are apparent between sites with great crested newt (GCN) absent (salmon) or present (green). Box and whisker plots show (a) SI2: pond area, (b) SI5: shade, (c) SI8: pond count, and (d) SI10: macrophyte scores for sites with GCNs present or absent. “n.s.” indicates statistically non-significant differences between SI scores in ponds with GCN absent or present; “**” indicates statistically significant differences at P<0.001 between SI scores in ponds with GCN absent or present.

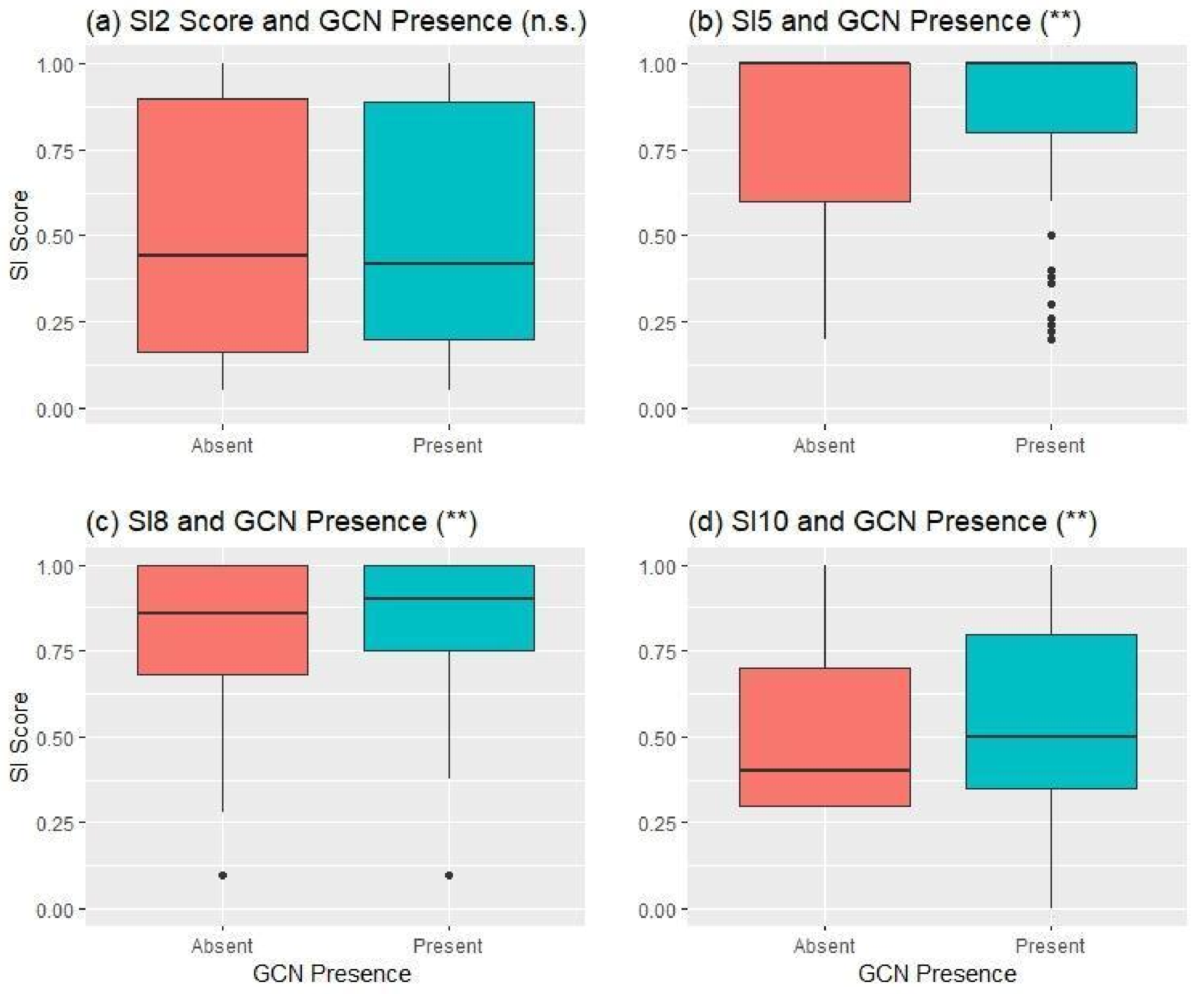

Figure 3: Most categorical suitability indexes (SIs; Table 10) for the great crested newt (GCN) show marginally higher frequency of ponds with GCN present (green) for higher score categories, and lower frequency of ponds with GCN absent (salmon) for lower score categories. This pattern is not apparent for waterfowl (SI6), and is hard to assess for pond location (SI1) due to the predominance of a score of 1, meaning few samples were from the edges of GCN distribution. Bar charts show the frequency of score levels for sites with GCNs present or absent for (a) SI1: geographic location, (b) SI3: pond permanence, (c) SI4: water quality, (d) SI6: waterfowl, (e) SI7: fish, and (f) SI9: terrestrial habitat. "n.s." indicates statistically non-significant differences between SI scores in ponds with GCN absent or present, "**" indicates the statistically significant differences at P<0.001 between SI scores in ponds with GCN absent or present.

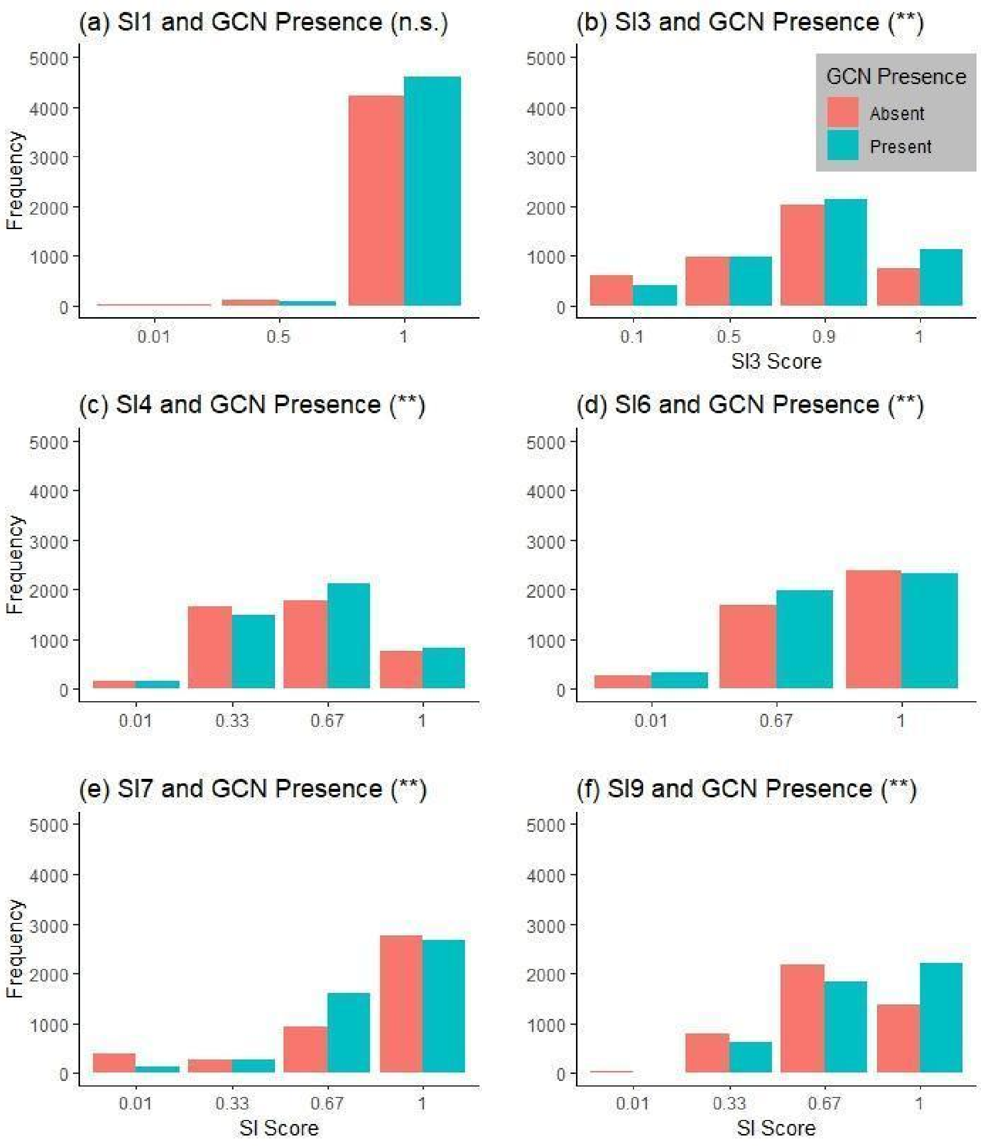

Figure 4:  Habitat suitability index (HSI) scores for the great crested newt (GCN) are higher in ponds where the species is present. HSI (a) categories and (b) scores are shown for sites with (salmon) or without (green) GCNs. (a) The bar chart shows higher frequencies of ponds with GCN present and lower frequencies of ponds with GCN present for the higher HSI categories (Average, Good, and Excellent). The panel also shows lower frequencies of ponds with GCN present and higher frequencies of ponds with GCN present for the lower HSI categories (Poor and Below average). (b) Box and whisker plot of scores showing higher HSI scores for sites where GCNs are present.

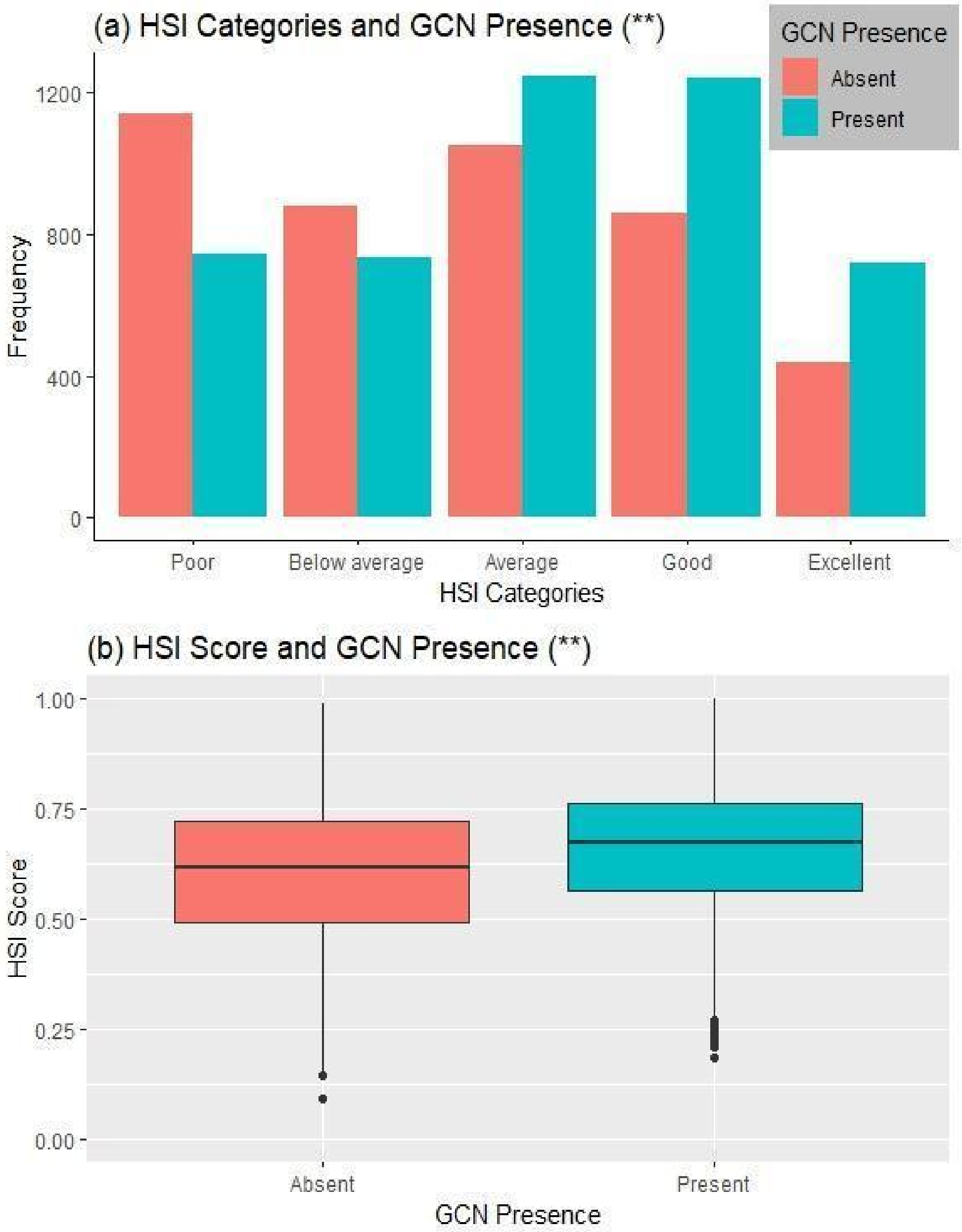

Figure 5: Distribution of (a) HSI categories and (b) HSI scores for sites with great crested newts (GCNs) present or absent under the newly proposed habitat suitability index (HSI). HSI scores are statistically higher in ponds where GCNs are present (P<0.001) - see Table 3 and Supp.Mat.3g. (a) Bar chart showing higher frequencies of ponds with GCN present and lower frequencies of ponds with GCN present for the higher HSI categories (Average, Good and Excellent). This panel shows lower frequencies of ponds with GCN present and higher frequencies of ponds with GCN absent for the lower HSI categories (Poor, Below average). (b) Box and whisker plot of the newly proposed HSI scores showing higher HSI scores for sites with where GCNs are present (P<0.001) - see Table 3 and Supp.Mat.3h. The interquartile range of modified HSI scores in Figure 5.B is higher and more compact than that of Oldham's HSI shown in Figure 4.B , necessitating the new category divisions proposed in Table 2.

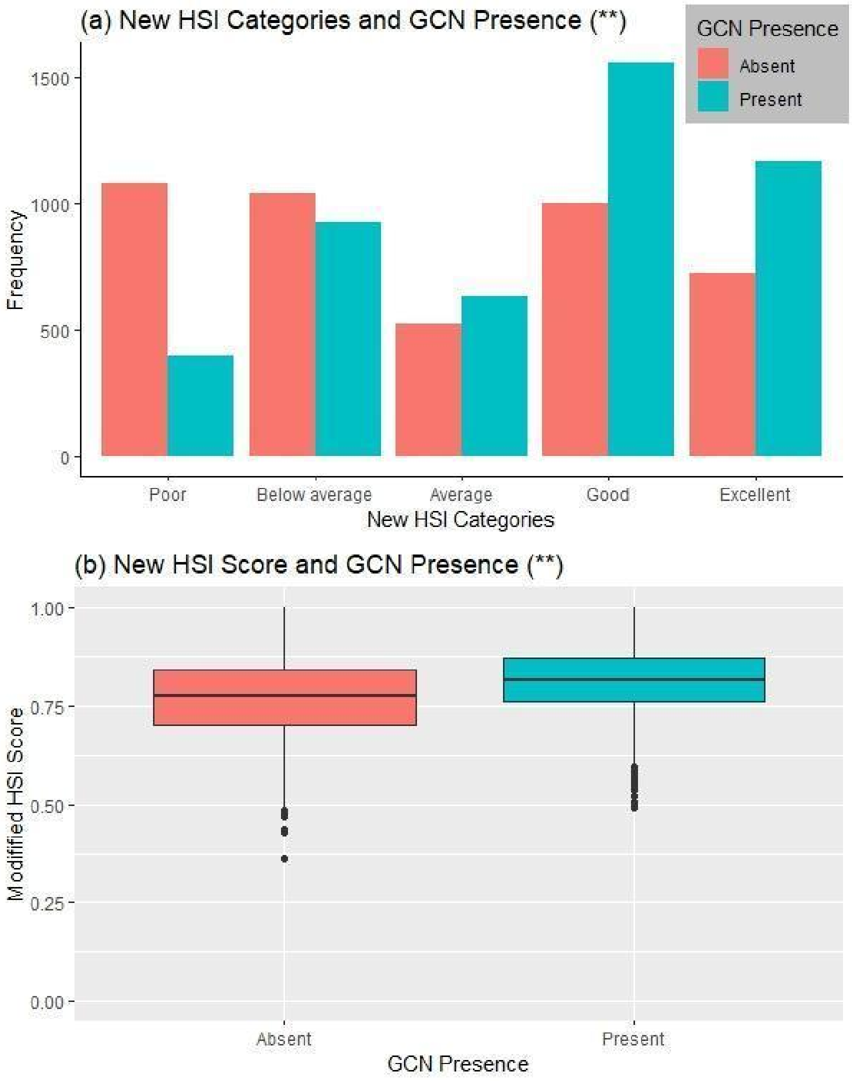

Figure 6. The newly proposed HSI outperforms the old HSI both in terms of effect sizes and goodness of fit when predicting presence/absence of the great created newt (GCN). Statistical results of the binomial regression testing the relationship between HSI scores and GCN presence/absence. The length of the arrows is proportional to the effect size for the output of the binomial regression, under the two HSIs, first tests on the original dataset of the paper, and secondly on the independent dataset used for verification. The circles show the $R^2$ values - with the new HSI, these values are larger (more so for the original dataset) suggesting a better goodness-of-fit between the new HSI and the GCN presence/absence data. The effect sizes are only statistically significant when using the larger, original dataset.

| | Original dataset - old HSI | Original dataset - new HSI | Verification dataset - old HSI | Verification dataset - new HSI |
|---|---|---|---|---|
| Effect size | ** 0.736 | ** 5.266 | 0.7759 | 1.411 |
| $R^2$ (calculated from glm as 1-(residual deviance/ null deviance) | 0.029 | 0.038 | 0.041 | 0.043 |

Notes:
**: P < 0.001, otherwise P >0.05.
Effect size: direction of arrow reflects direction of effect, length of arrow reflects size of effect.
$R^2$: Radius of circle reflects size of $R^2$ value.
