## Supplementary material for "An improved habitat suitability index for the great crested newt": Tables

Table 1: Summary of suitability indices from the great crested newt habitat suitability index, and their corresponding environmental criteria. This is a summary of the information provided in Oldham et al. (2001) and ARG UK (2010).

| **Suitability Index** | **Environmental criteria** | **Details** |
|---|---|---|
| SI1 | Geographic location | 3 categories giving scores of 0.01, 0.5 or 1. |
| SI2 | Pond area | Graph gives scores for pond areas measured in $m^2$. |
| SI3 | Pond permanence | 4 categories giving scores 0.1-1. |
| SI4 | Water quality | 4 categories giving scores 0.01, 0.33, 0.67 or 1, based on invertebrate diversity |
| SI5 | Shoreline shade | Graph gives scores for percentage of shoreline shaded. |
| SI6 | Waterfowl | 3 categories giving scores of 0.01, 0.67 or 1. |
| SI7 | Fish | 4 categories giving scores 0.01, 0.33, 0.67 or 1. |
| SI8 | Pond density | Graph gives scores using the number of ponds within 1km divided by $\pi$. |
| SI9 | Terrestrial habitat | 4 categories giving scores 0.01, 0.33, 0.67 or 1. |
| SI10 | Macrophyte cover | Graph gives scores for percentage of surface area covered by macrophytes. |

Table 2: Potential modifications to improve the great crested newt (GCN) suitability indices (SIs) within the habitat suitability index (HSI), inferred from peer-review literature, user survey results, and ecological data analysis performed in this research.

| **Aspect of the HSI** | **Potential Modifications to HSI or associated guidance** |
|---|---|
| SI1 | Increase the number of zones, use more up-to-date data, and better define borders. Include new map with new Scotland zones (O'Brien et al., 2017). Make an easier-to-read map or an online look-up system |
| SI2 | Create SI2 scores for ponds over 2,000 $m^2$. Removing or rescoring SI2 based on underlying environmental data (Denoel & Ficetola, 2008). |
| SI3 | Test relationship between GCN presence/absence and desiccation (Griffiths & Williams, 2000). |
| SI4 | Removing SI4 from HSI. Replace with chemical testing of water. |
| SI6 | Removing SI6 from HSI. |
| SI7 | Test importance of fish presence for GCN; try removing from HSI. |
| SI8 | Trial exclusion of SI8 from HSI if HSI <0.75 (Oldham et al., 2000). Ensure SI8 scores correctly divided pond numbers by pi; create new graph to avoid need to divide by pi. Test importance of pond density for GCN. Removing from HSI |
| SI10 | Change focus from living plant material to any egg-laying material. |

Table 3: Potential modifications to improve the great crested newt (GCN) the habitat suitability index (HSI), inferred from peer-review literature, user survey results, and ecological data analysis performed in this research.

| |
|---|
| Improve guidance to reduce subjectivity and emphasise appropriate use (not as substitute for population survey). Buxton et al., 2021. |
| Weight SIs according to importance (Oldham et al., 2000). |
| Add measures of uncertainty to output. Bender et al., 1996; Burgman et al., 2001; Zajac et al., 2015; Biggs et al., 2014. |
| Test predictive ability of HSI scores on GCN presence/absence, abundance, and breeding success. Reason et al., 2013; Buxton et al., 2021; Lewis et al., 2007. |
| Create clearer graphs and provide formulae for the relationship between the SI score and the underlying environmental variable shown in the graph. |
| Check if HSI scores provided correctly uses the geometric mean of SIs; try arithmetic mean or cumulative score of SIs to reduce mistakes of miscalculation. U.S. Fish and Wildlife Service, 1981; Burgman et al., 2001. |
| Increase the number of categories for categorical SIs. |
| Investigate additional factors for potential inclusion (Langton et al., 2001; Skei et al., 2006; Marklund et al., 2002; Denoel & Ficetola, 2008; Gustafson et al., 2011; Cresswell & Whitworth 2004; JNCC, 2019; Dunford & Berry, 2013. |
| Provide greater clarity on detailed, reproducible methods for recording SIs. |

Table 4: Great crested newt habitat suitability index (HSI) categories (left column) and corresponding values (centre column) under the original habitat suitability index (ARG UK, 2010) and with the new modified habitat suitability index (right column).

| **Category** | **HSI value cut-offs for original HSI, as given in ARG (2010)** | **HSI value (x) cut offs for new HSI combining useful modifications** |
|---|---|---|
| Poor | <0.5 | x <0.70 |
| Below Average | 0.5-0.59 | 0.70 ≥ x < 0.77 |
| Average | 0.6-0.69 | 0.77 ≥ x < 0.80 |
| Good | 0.7-0.79 | 0.80 ≥ x < 0.87 |
| Excellent | >0.8 | x ≥0.87 |
